# The pioneer transcription factor Zelda facilitates the exit from regeneration and restoration of patterning in *Drosophila*

**DOI:** 10.1101/2024.05.30.596672

**Authors:** Anish Bose, Keaton Schuster, Chandril Kodali, Surabhi Sonam, Rachel Smith-Bolton

## Abstract

For a damaged tissue to regenerate, the injured site must repair the wound, proliferate, and restore the correct patterning and cell types. We found that Zelda, a pioneer transcription factor largely known for its role in embryonic zygotic genome activation, is dispensable for normal wing development but crucial for wing disc patterning during regeneration. Impairing Zelda function during disc regeneration resulted in adult wings with a plethora of cell fate errors, affecting the veins, margins, and posterior compartment identity. Using CUT&RUN, we identified and validated targets of Zelda including the cell fate genes *cut, Delta* and *achaete*, which failed to return to their normal expression patterns upon loss of Zelda. In addition, Zelda controls expression of factors previously established to preserve cell fate during regeneration like *taranis* and *osa,* which stabilizes *engrailed* expression during regeneration, thereby preserving posterior identity. Finally, Zelda ensures proper expression of the integrins encoded by *multiple edematous wings* and *myospheroid* during regeneration to prevent blisters in the resuting adult wing. Thus, Zelda is crucial for maintaining cell fate and structural architecture of the regenerating tissue.

## Introduction

The phenomenon of regeneration after damage or loss of a limb, organ, or tissue involves regrowth of the tissue, followed by cell fate re-specification and differentiation to restore the original structure and function. Some animals like axolotls have incredible regenerative capacity and can restore lost limbs and transected spinal cords (Poss, 2010). However, in most mammals, while some tissues and organs such as the liver, skeletal muscles, and skin can regenerate well, their regenerative potential reduces with age (Yun, 2015), and tissues like limbs, joints, and the heart regenerate very poorly throughout their lifespan (Mokalled & Poss, 2018; Rafii et al., 2016; Wells & Watt, 2018). Thus, understanding the damage-response mechanisms in organs and appendages that are capable of regeneration can lead to a broader understanding of how the regeneration process occurs in different contexts and how it may be applied to induce wound healing and repair. Importantly, while numerous studies have focused on how regeneration begins and progresses in a variety of model organisms, little is understood about how regeneration ends and normal cell fate and function are restored.

The *Drosophila* wing imaginal disc, the larval precursor of the adult wing, for example, has the intrinsic ability to regenerate (reviewed in Hariharan & Serras, 2017). During wing disc regeneration, the damaged tissue proliferates and organizes cell fates to re-pattern the tissue and form the adult wing (Hariharan & Serras, 2017). Reports over the last decade have shown that wing disc regeneration requires regeneration-specific factors to regulate cell fate as the tissue repairs itself (Abidi et al., 2023; Schuster & Smith-Bolton, 2015; Tian & Smith-Bolton, 2021). However, the regulatory mechanisms that control cell fate during wing regeneration remain poorly understood, raising the question: how does the regenerating tissue establish cell fate and patterning in the damaged wing disc as it exits regeneration and returns to normal development?

Here, we show that the pioneer transcription factor *zelda* (*zld*) has a regeneration-specific function in wing imaginal discs. Zld is essential during embryogenesis for the maternal-to-zygotic transition, pattern formation, sex determination, and cellularization (Harrison et al., 2011; Liang et al., 2008; McDaniel et al., 2019; Nien et al., 2011; Schulz et al., 2015; Staudt et al., 2006; Sun et al., 2015b). Outside of the embryo, zygotic *zld* is important for maintaining type II neuroblasts in an undifferentiated state in the fly brain (Larson et al., 2021). However, very little is known about the role of *zld* during wing development.

In this study, we show that *zld* is dispensable for normal wing development but is required during wing imaginal disc regeneration. Specifically, we found that *zld* is important for controlling tissue patterning and cell fate specification through a combination of regulating specific cell fate determinants as well as ensuring a timely transition from regeneration gene expression to developmental gene expression. We found that reducing or impairing Zld disrupted margin and vein cell fate, induced posterior-to-anterior fate transitions, and interfered with integrin expression, resulting in adult wings with blisters. Thus, *zld* is crucial for re-establishing and stabilizing cell fate, regulating patterning, and ensuring correct wing architecture, allowing a regenerated wing disc to undergo metamorphosis to generate the adult wing blade.

### A spatio-temporal system to cause tissue damage in the wing imaginal disc

To explore how cell fate and pattern are re-introduced in late regeneration and identify the mechanisms that control gene expression during this phase of repair, we used a spatio-temporal damage system to induce damage and regeneration in the *Drosophila* wing imaginal disc (Brock et al., 2017; Smith-Bolton et al., 2009). This method uses a GAL4 transgene inserted in the *rotund* (*rn*) locus (St Pierre et al., 2002), which is expressed in the pouch of wing discs, the primordium of the adult wing blade (Figure 1A). GAL4 activity is suppressed at 18°C by a temperature-sensitive Gal80 (Gal80^ts^) (McGuire et al., 2003). A temperature shift to 30°C in the early third instar (day 7 after egg lay) allows the GAL4 to express the pro-apoptotic *UAS-reaper* (*rpr*) (Aplin & Kaufman, 1997) to induce cell death. After 24 hours, shifting the temperature back to 18°C allows Gal80^ts^ inhibition of *rpr* expression (Figure 1A). The regenerating wing pouch, marked by expression of *nubbin* (*nub*), had very few cells remaining at 0 hours of recovery time (R0). The wing pouch steadily increased in size from 24 hours (R24) through 48 hours (R48) and was almost back to its normal size by 72 hours (R72) (Figure 1A).

**Figure 1:**
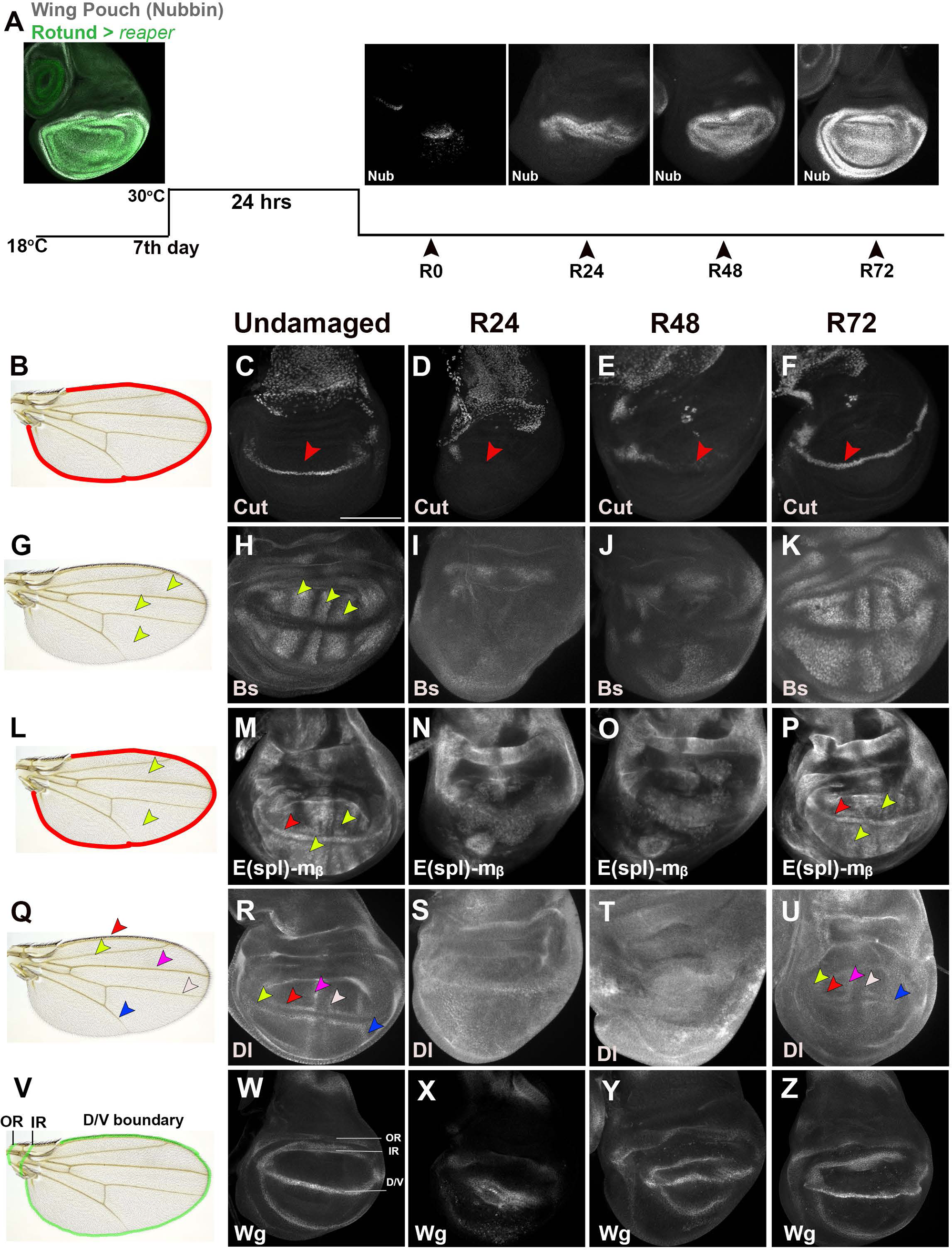
Gene expression during regeneration. A) Wing pouch of 3rd-instar wing discs with immunostaining for Nubbin (gray) and rn>GFP (green) demarcating where *rpr* was expressed. B) Adult wing with margin outlined in red. C-F) Ct expression at the D/V boundary (red arrowheads) in undamaged (C), R24 (D), R48 (E), and R72 (F) wing discs. G) Adult wing with some intervein regions marked by yellow arrowheads. H-K) Bs-GFP expression in the intervein regions (yellow arrowheads) in undamaged (H), R24 (I), R48 (J), and R72 (K) wing discs. L) Adult wing with the margin outlined in red and intervein regions marked with yellow arrowheads. M-P*) E(spl)Mb-CD2 (CD2)* expression in the margin (red arrowhead) and intervein regions (yellow arrowheads) in undamaged (M), R24 (N), R48 (O), and R72 (P) wing discs. Q) Adult wing with veins marked by arrowheads: L1 (red), L2 (yellow), L3 (magenta), L4 (white), and L5 (blue). R-U) Dl immunostaining in pro-vein cells with arrowheads the same colors as in (Q) in undamaged (R), R24 (S), R48 (T), and R72 (U) wing discs. V) Adult wing with green lines marking the D/V boundary and approximate locations of the Wg IR and OR. W-Z) Wg immunostaining in undamaged (W), R24 (X), R48 (Y), and R72 (Z) wing discs. Scale bar: 100µM.

Several such methods of inducing damage have been used to identify changes in patterning and cell fate that occur in the regenerating wing disc (Díaz-García & Baonza, 2013; Repiso et al., 2013; Smith-Bolton et al., 2009). The regenerating pouch loses expression of markers for various cell fates, which are not re-established until the end of regeneration (Díaz-García & Baonza, 2013; Smith-Bolton et al., 2009). For example, the margins of the adult wing (Figure 1B) are established by expression of the gene *cut* (*ct)* at the dorso-ventral (D/V) boundary of the wing disc downstream of Notch signaling (Figure 1C) (Jack et al., 1991; Krupp et al., 2005). After ablation, Ct expression was absent at R24 (Figure 1D) even though the D/V boundary and Notch signaling were present (Smith-Bolton et al., 2009). Ct expression partially returned at R48 (Figure 1E) and was completely restored by R72 (Figure 1F).

In addition, the regenerating tissue loses intervein and provein marker expression (Díaz-García & Baonza, 2013; Repiso et al., 2013; Smith-Bolton et al., 2009). The intervein regions between the veins in the adult wing can be identified in the developing disc by expression of *blistered* (*bs*) (Nussbaumer et al., 2000; Roch et al., 1998), which was absent in the pouch at R24, was mostly absent in the pouch at R48 except for a few Bs-positive cells at the edges of the pouch, and was finally restored at R72 in 5/8 wing discs (Figure 1G-K). Vein and intervein gene expression is regulated by Notch (N) signaling activity (de Celis et al., 1997), which we detected using the *E(spl)Mb-CD2* N signaling reporter (Panin et al., 1997), and is expressed at the D/V boundary and in the intervein regions during normal development (Figure 1L-M). At R24, the N activity reporter was expressed in a thick stripe around the D/V boundary, which persisted at R48 (Figure 1N-O). By R72, expression was restored to the D/V boundary and intervein regions in 3/5 discs (Figure 1P), which is similar to previous reports (Smith-Bolton et al., 2009). We assessed the pro-vein cell identity (Figure 1Q) using expression of the N signaling ligand *Delta (Dl)* (Figure 1R) (Díaz-García & Baonza, 2013; Doherty et al., 1996). At R24, Dl was expressed in a thin stripe at the presumptive D/V boundary (Figure 1S), which becomes broader and more diffuse by R48 (Figure 1T). Normal Dl expression returned by R72 in 4/10 discs (Figure 1U), while the remaining discs had incomplete Dl expression that was missing sections such as the L2 and L5 proveins.

The WNT family ligand *wingless* (*wg*) is essential for wing blade, margin and hinge formation (Baker, 1988; Rosales-Vega et al., 2023; Williams et al., 1993) (Figure 1V), and is expressed along the D/V boundary and in an inner ring (IR) and an outer ring (OR) around the wing pouch (Rodríguez et al., 2002) (Figure 1V-W). At R24, Wg was expressed throughout the regenerating blastema, as previously reported (Smith-Bolton et al., 2009) (Figure 1X). The Wg expression pattern began transitioning into the D/V boundary and hinge rings at R48, passing through intermediate expression patterns not observed during normal development, and by R72 returned to its normal pattern (Figure 1Y-Z). Thus, cell fate genes progress through non-developmental expression patterns during regeneration before returning to their correct expression patterns. As noted, not all regenerating discs had completely normal cell fate gene expression by R72, which was reflected in the resulting adult wings that had minor errors in patterning (Abidi et al., 2023; Schuster & Smith-Bolton, 2015; Tian & Smith-Bolton, 2021).

### Zelda regulates cell fate and patterning during late regeneration

We wondered what factors facilitate the shift back to normal development by three days after damage. We hypothesized that this exit from regeneration could be orchestrated by a transcription factor that facilitates transitions between developmental states, such as the pioneer transcription factor Zelda (Zld).

Previous work has suggested that the *zld* transcript is not uniformly expressed in wing discs (Staudt et al., 2006). However, our immunostaining showed Zld protein is expressed uniformly throughout the developing disc (Figure 2A-A”). At R24, Zld expression was slightly but not statistically significantly elevated in the regenerating pouch (Figure 2B-B”). Zld expression was markedly increased at R48 within the regenerating region and remained upregulated at R72 (Figure 2C-E). We also observed a similar upregulation in *zld* mRNA as assessed by qPCR (Figure 2F).

**Figure 2:**
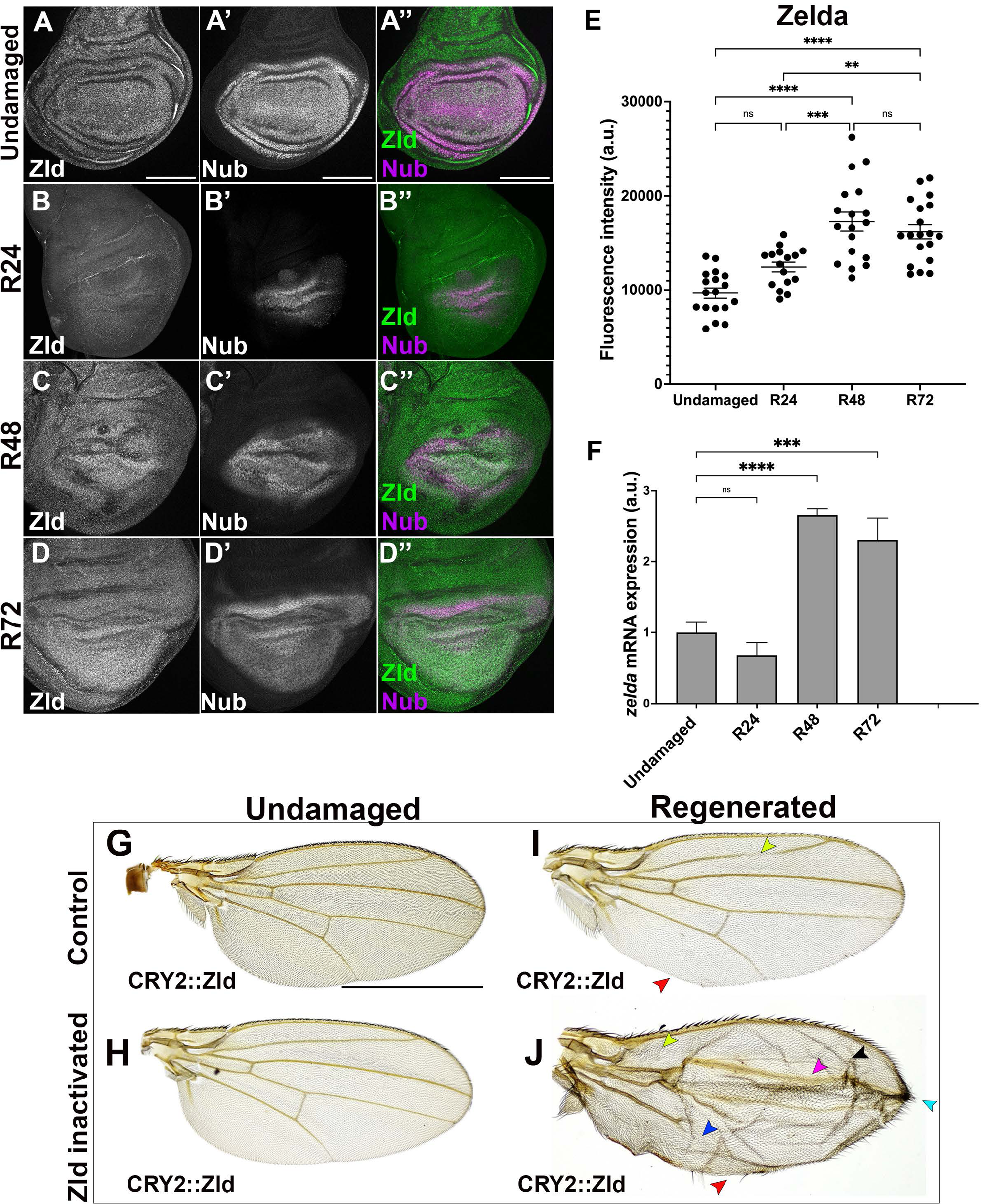
Zld is important for cell fate and patterning during regeneration. A-D) Undamaged, R24, R48 and R72 *w^1118^* wing discs immunostained for Zld (A-D), Nubbin (A’-D’). Merged (A”-D”). E) Quantification of Zld immunostaining fluorescence intensity. n=20 discs each. ns> 0.05, **p< 0.01, ***p< 0.001, ****p< 0.0001, One-way ANOVA test. F) Quantification of *zld* mRNA by qPCR. One-way ANOVA test, ***p<0.001, ****p<0.0001. G) Adult wing from a *CRY2::zld* fly not exposed to blue light. H) Adult wing from a *zld::CRY2* fly without disc damage exposed to blue light between R0 and R72. I) Adult wing after disc regeneration in a *CRY2::zld* fly not exposed to blue light having some defects like incomplete L2 vein (yellow arrowhead) and incomplete posterior margin (red arrowhead). J) Adult wing after disc regeneration in a *CRY2::zld* fly exposed to blue light between R0 and R72 having missing L2 vein (yellow arrowhead), altered L3 vein thickness (magenta arrowhead), incomplete L5 vein (blue arrowhead), incomplete posterior margin (red arrowhead), blister formation (black arrowhead) and distal edge vein material (teal arrowhead). Wing disc scale bars: 100µm. Adult wing scale bar: 750µm.

To determine Zld’s role during regeneration, we used a previously reported fly line in which the endogenous Zld is tagged with cryptochrome 2 (CRY2) at its N terminus (CRY2::Zld) to optogenetically inactivate Zelda (McDaniel et al., 2019). We inactivated Zld by shining blue LED light on the larvae during R0-R72. Exposing undamaged *w^1118^* larvae that did not contain the *CRY2::zld* to acute LED blue light led to adult wings that were normal (Figure S1A,C). Exposing regenerating discs in *w^1118^* larvae that did not contain the *CRY2::zld* to acute LED blue light led to 50% (n=98) of adult wings with minor defects, which was similar to the 58% (n=92) of adult wings with minor defects for *w^1118^* animals that were not exposed to blue light during disc regeneration, indicating that blue light exposure does not cause additional defects during development or regeneration (Figure S1B,D). In addition, the CRY2 insertion does not interfere with normal wing development (Figure 2G) or disc regeneration, with sporadic minor defects in adult wings similar to controls, where 56% (n=230) (Figures 2I, S1B) of adult wings after regeneration had defects. Inactivation of CRY2::Zld by blue light during the third instar did not cause any defects in the adult wing (Figure 2H), suggesting that Zld is not required for normal wing development.

Interestingly, exposing *CRY2::zld* larvae with damaged discs to blue light between R0 and R72 led to 96.8% (n=218) of adult wings with defects. Inactivation of Zld during disc regeneration resulted in numerous defects, such as missing or incomplete L2 and L5 veins, altered L3 and L4 vein thickness, missing cross-veins, and ectopic vein material at the distal end of the wing. A percentage of each margin was also missing, especially in the posterior portion of the wing. Finally, these adult wings had a high rate of blisters (Figure 2J). Additionally, we found some variability in the frequency of these defects, likely due to the larvae positioned at different depths in the food, resulting in inequal exposure to blue light. However, the phenotypes observed were consistent and reproducible.

To validate that these defects were due to loss of Zld function, we built a *zld* RNAi transgene by placing a previously published *zld*-specific RNAi sequence (Sun et al., 2015) under the control of a LexA operator (LexAop) (Chang et al., 2022) to avoid regulation by the Gal80^ts^. The *LexAop zld-RNAi* was expressed using a *pdm2-LexA* driver that is expressed in the wing disc pouch and hinge region (Figure S1E). Expression of the *LexAop zld-RNAi* during normal development in the wing disc reduced Zld levels (Figure S1F-G”) but did not lead to any defects in the adult wing, again indicating that that Zld is not required for normal wing development (Figure 2N). Expression of *zld*-*RNAi* also reduced Zld levels at R24 (Figure S1H-I”), R48 (Figure S1J-K”), and R72 (Figure S1L-M”). This reduction yielded adult wings with similar defects to the CRY2::Zld system, such as missing and/or incomplete veins, ectopic vein material at the distal edge of the wing, missing margins, and blisters (Figure S1Q-S). Adult wings from *pdm2-LexA*; *attP2* damaged discs in most cases appeared well formed, with 65% of wings regenerating perfectly (n=131) (Figure S1P). Regenerating discs containing only the *LexAop-zldRNAi* transgene resulted in adult wings with mild defects that were nevertheless similar to those seen with Zld knockdown during regeneration, suggesting that this *LexAop* promoter might allow leaky expression of the RNAi (Figure S1O). Thus, inactivation of Zld by CRY2 and RNAi-mediated knockdown of Zld after damage resulted in many patterning and cell fate defects, as well as blistering in the adult wing.

### Zld is enriched at developmental genes during late regeneration

To identify the genes Zld regulates to facilitate correct regeneration, we performed the protein-DNA binding assay <u>C</u>leavage <u>U</u>nder <u>T</u>argets and <u>R</u>elease <u>U</u>sing <u>N</u>uclease (CUT&RUN) (Skene & Henikoff, 2017) to find where Zld binds throughout the *Drosophila* genome in undamaged and regenerating wing discs. Our CUT&RUN data showed that Zld had very few enriched binding sites near transcription start sites (TSSs) in undamaged wing discs (Figure 3A), supporting our conclusion that Zld is dispensable for normal wing development. However, we found that Zld bound to thousands of genomic loci in late regenerating wing discs that are near TSSs (Figure 3A). We compared the genomic loci bound by Zld during regeneration with previously published ChIP-seq data for Zld during stage 14 of embryonic development (Harrison et al., 2011) and in Type II neuroblasts of the larval brain (Larson et al., 2021). Sixty-nine percent of the loci bound by Zld during wing regeneration were also found in ChIP-seq datasets for stage 14 embryos, while 31% of the loci bound by Zld during wing regeneration were also bound by Zld in Type II Neuroblasts (Figure S2A). Further analysis showed that Zld binding was enriched within 1kb of gene promoters (Figure 3B), where 50% of Zld-bound sites were located (Figure 3B). When we compared this distribution of binding sites to Zld’s binding profiles in the embryo and the brain, we found that the tissues show different profiles (Figure S2B-D).

**Figure 3:**
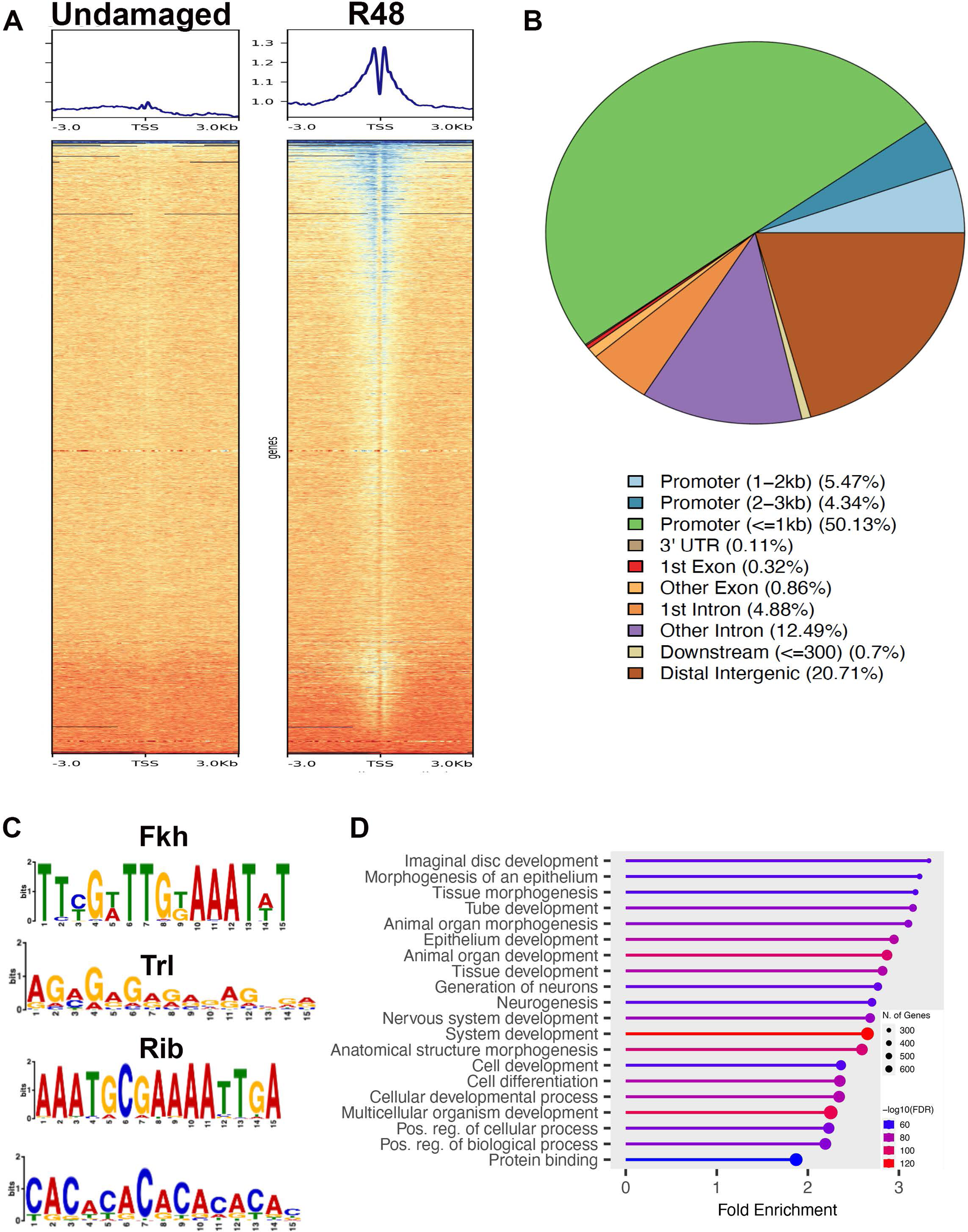
Zld binds near genes important for wing development and morphogenesis. A) Heatmap of Zld occupancy in undamaged and regenerating wing discs +/− 3kb from Transcription Start Sites (TSSs). B) Location of Zld binding sites relative to genes. C) Motif discovery and enrichment analysis of sequences bound by Zld at R48. Enriched motifs were Fkh (MEME, E-value: 1.5e-086), Trl (STREME, E-value: 2.7e-002), Rib (MEME, E-value: 1.3e-039), and CA-rich repeats (MEME, E-value: 3.1e-063). D) Biological processes enriched in the genes near Zld binding sites according to GO analysis.

We used the sequences derived from the Zld binding sites during wing regeneration to find enriched motifs that Zld or its cofactors might bind to. The pioneering activity of Zld in early embryos is due to its ability to bind to CAGGTA sequences throughout the genome (Harrison et al., 2011), but Zld binds other sequences in transcriptionally active tissues such as the type II neuroblasts. (Larson et al., 2021). One common co-factor for Zld in both tissues is GAF/Trithorax-like (Trl), which recognizes GA-rich repeats (Gaskill et al., 2021; Larson et al., 2021). Zld binding sites in regenerating wing discs at R48 were not enriched for CAGGTA sites, but did contain GA-rich repeats, suggesting that Trl/GAF could be acting as a Zld cofactor during wing disc regeneration (Figure 3C). Zld binding sites were also enriched for motifs that are recognized by other transcription factors such as Forkhead (Fkh) and Ribbon (Rib), and the importance of these transcription factors for wing disc regeneration will be of interest in future studies (Figure 3C). Thus, our motif discovery analysis suggests that Zld may work with the known cofactor Trl/GAF and may also work with other partners during late regeneration in wing imaginal discs.

We assessed the list of genes near Zld binding sites in regenerating wing discs using Gene Ontology (GO) tools. Interestingly, we found enrichment for genes involved in various developmental and morphogenesis processes (Figure 3D). To validate our CUT&RUN data, we assessed expression of genes near putative Zld binding sites individually. One such gene was *fz2*, which encodes a receptor for WNT family ligands such as Wg and is important for developmental processes dependent on WNT signaling (Chaudhary et al., 2019). Fz2 is expressed in the notum as well as the dorsal and ventral regions of the wing pouch (Figure S2F). Inactivation of Zld during normal development did not change Fz2 expression levels or patterning (Figure S2G,J). During late regeneration, Fz2 was expressed throughout the regenerating pouch at R72 (Figure S2H) and was markedly reduced upon inactivation of Zld (Figure S2I,J). Thus, we established that a developmental patterning gene near a Zld binding site requires Zld for full expression during regeneration. Hence, we wondered if Zld could be important for correct expression of other developmental genes during regeneration.

### Zld regulates genes important for wing margin and sensory bristle fate

Adult wings that developed from damaged and regenerated discs in which Zld had been inactivated or knocked down had very specific defects in veins, margins, and sensory bristles. Hence, we wondered if Zld could be responsible for inducing expression of specific genes required for these cell fates. First, we confirmed that inactivation or knockdown of Zld during normal development did not lead to any vein disruption in adult wings defects in the wing margin, or loss of sensory bristles on the anterior margin (Figures 4A-B, S3A). Adult wings formed from control wing discs that had the CRY2 insertion but regenerated without blue light exposure or had the genotype *pdm2-LexA/+; attP2/+,* were well patterned, with occasional disruption of veins and margins (Figures 4C, S3B).

**Figure 4:**
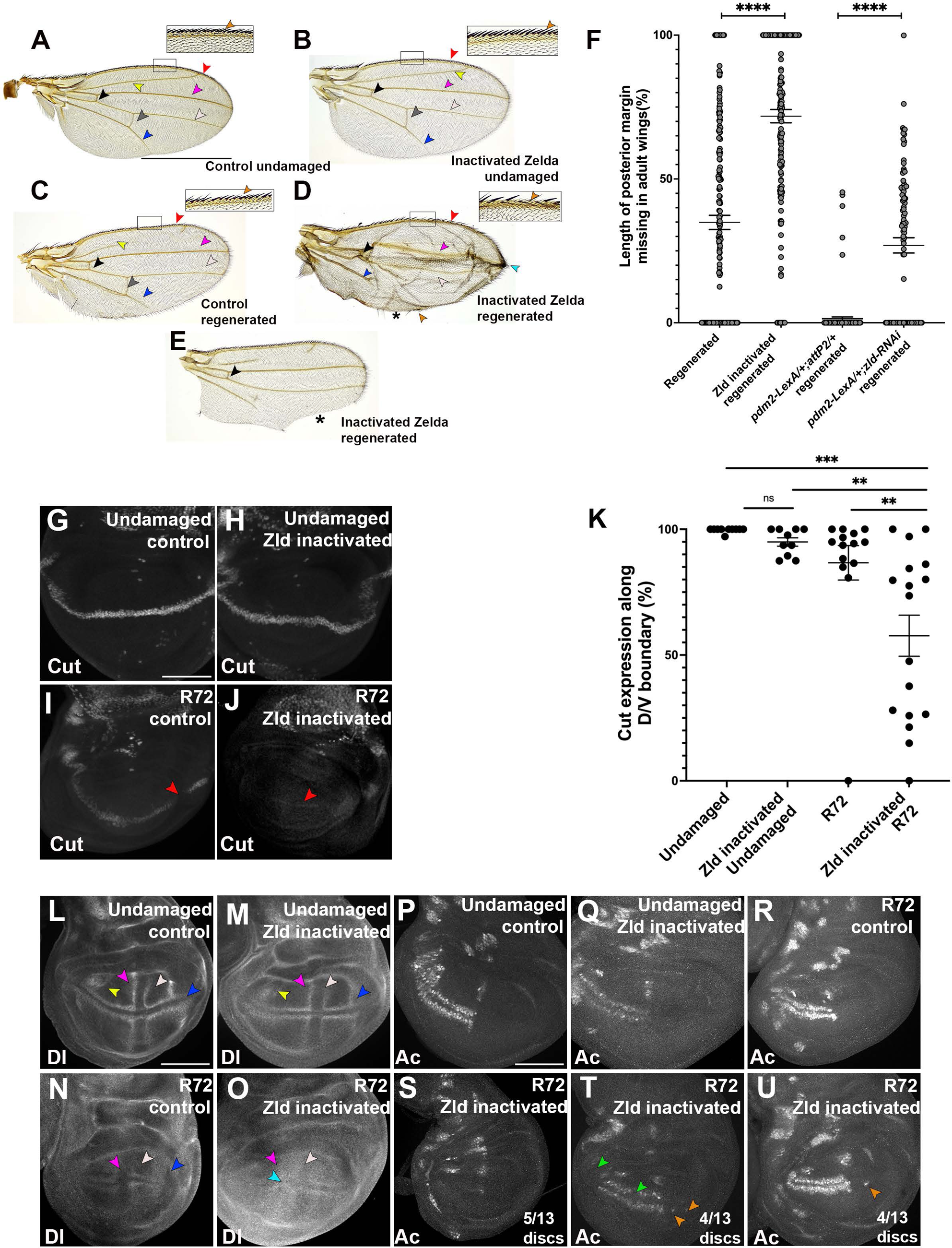
Zld is important for margin, vein, and sensory organ fate after regeneration. A) Undamaged adult wing showing ACV (black), PCV (Grey), L1 (red), L2 (yellow), L3 (magenta), L4 (white), L5 (blue) veins and sensory bristles (orange, outlined box with higher magnification inset). B) Adult wing with Zld inactivated during normal development, arrowheads same as in (A). C) Adult wing after disc regeneration, arrowheads same as in (A). D-E) Adult wings after Zld inactivation during regeneration, arrowheads same as in (A). * denotes missing posterior margin. F) Quantification of missing posterior margin for control wings after regeneration (n=207), Zld inactivated wings after regeneration (n=190), control *pdm2-LexA;attP2/+* RNAi wings after regeneration (n=131) and pdm2-LexA;zld-RNAi/+ wings after regeneration (n=91), ****p<0.0001, students t-test. G-K) Ct immunostaining in an undamaged wing disc (G), undamaged disc with Zld inactivated (H), R72 wing disc (I), R72 wing disc with Zld inactivated (J) and quantification of Cut expression along D/V boundary for undamaged discs: control (n=10), Zld inactivated (n=10) and R72 discs for control (n=14) and Zld inactivated (n=17). ns p >0.5, **p<.01, ***p<0.001, One-way ANOVA (K). L-O) Dl immunostaining in an undamaged wing disc (L), undamaged wing disc with Zld inactivated (M), R72 regenerating wing disc (N), and R72 regenerating wing disc with Zld inactivated (O). Arrowheads mark L2 (yellow), L3 (magenta), L4 (white), and L5 (blue). P-U) Ac immunostaining in an undamaged wing disc (P), undamaged wing disc with Zld inactivated (Q), R72 wing disc (R), and R72 wing disc with Zld inactivated (S-U). Scale bar for adult wings: 750µM. Scale bar for all discs: 100µM.

Finally, we analyzed adult wings that arose from wing discs that regenerated while Zelda was inactivated or knocked down. A high percentage of these wings had errors including poorly defined L3 and L4 veins and margin missing in sections along with absent sensory bristles on the anterior side (Figures 4D, S3C). In addition, large sections of the L2 and L5 veins were missing (Figures 4D-E, S3D-E), and anterior and posterior crossveins were missing (Figure S3F-G).

These wings also had aberrant vein tissue on the distal edge of the adult wing (Figures 4D, S3H) and had substantial sections of margin missing on the posterior edge (Figure 4E-F). Thus, Zld appears to be crucial for vein and margin cell fate specification during disc regeneration.

To understand how Zld is regulating these cell fates, we asked if Zld binds near genes important for margin and vein identity using our CUT&RUN data. First, we looked for genes near Zld binding sites that are important for margin fate. Interestingly, at R48, Zld is bound at the promoter region of *ct* (Figure S3I), which is expressed in the D/V boundary of developing wing discs and is important for margin fate (Jack et al., 1991; Krupp et al., 2005; Micchelli Craig A. et al., 1997) (Figure 1B-C). Ct expression along the D/V boundary remained stable upon Zld inactivation during normal development, as expected (Figure 4G-H). As previously shown, Ct expression was lost during regeneration, and re-appeared by R72 (Figure 1B-F). On average, Ct was expressed in 87% of the DV boundary by R72 (Figure 4I,K), indicating restoration of most of the margin. The gaps in Ct expression were likely responsible for the sporadic sections of margin missing in the resulting adult wings (Figure 4F). After Zld inactivation, R72 discs had variable expression of Ct, with some discs missing over 90% of Ct expression (Figure 4J). On average, only 57% of the D/V boundary expressed Ct by R72 (Figure 4K), which correlated with the high percentage of the adult wing margin missing after regeneration in wing discs lacking functional Zld.

We also looked for genes near Zld binding sites that are important for vein identity. Zld bound at the promoter as well as intronic regions of Dl (Figure S3J), which is expressed in the provein cells in developing wing discs and is important for provein fate and consequently intervein fate (de Celis et al., 1997; Huppert et al., 1997). There was no difference in Dl expression between undamaged discs with and without functional Zld (Figure 4L-M). As previously shown, Dl expression transitioned from a thin stripe at R24 to a broader and more diffused signal at R48 (Figure 1S-T). However, upon Zld inactivation, R48 discs had a markedly reduced Dl signal (Figure S3K-M), suggesting that Dl is a target of Zld. Furthermore, while Dl expression in control regenerating discs at R72 had mostly returned to normal, with L2 and some L5 provein cells missing Dl expression in a few discs (Figure 4N), 64% (n=14) of R72 discs that lacked Zld fucntion had incomplete expression of Dl (Figure 4O). The errors included L2 and L5 proveins lacking expression and weak expression in the L3 and L4 proveins (Figure 4O). Dl was also expressed diffusely in between the presumptive L3 and L4 proveins (Figure 4O), which is the region that gives rise to the tip of the wing and thus could be responsible for the extra vein material on the distal edge of the wing (Figures 4D, S1Q-S and S3C).

Tracts of sensory bristles were also often missing along the anterior margin of the wing after disc regeneration without Zelda (Figure 4A-D). Expression of the transcription factor *achaete* (*ac*) is crucial for sensory bristle development (Couso et al., 1994) and was identified as a putative target of Zld at R48 in our CUT&RUN analysis (Figure S3N). During normal development, Ac-expressing cells are found in the anterior half of the wing disc flanking the D/V boundary (Figure 4P) and are unaffected upon Zld inactivation (Figure 4Q). In R72 regenerating discs, Ac-expressing cells were present as expected (Figure 5R). However, in R72 discs that lacked Zld (Figure 4S-U), 38% had few Ac+ cells (Figure 4S), 31% had a reduced number of Ac+ cells (Figure 4T), and 31% had evenly distributed Ac+ cells in the anterior compartment (Figure 4U). This loss of Ac correlated with absent sensory bristles in the adult wing. Interestingly, we saw a percentage of wings with aberrant Ac expression on the posterior side of the wing disc (Figure 4T-U), suggesting Zld may play a role in maintaining proper posterior fate during regeneration. Taken together, our results show that Zld is important for the re-establishment of margin, sensory bristle, and vein fates in the regenerating wing disc, by regulating expression of targets like *ct*, *ac,* and *Dl* respectively.

**Figure 5:**
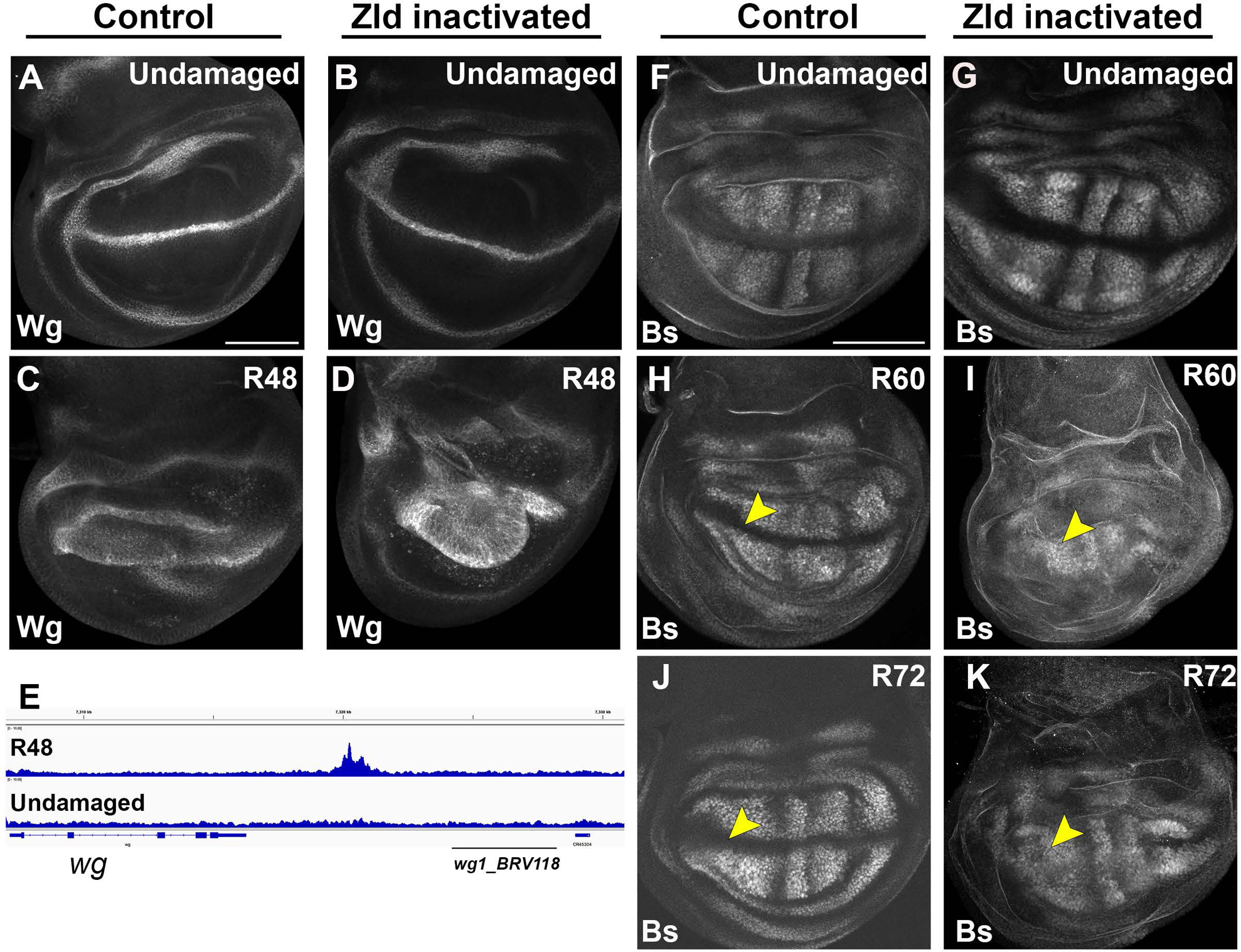
Zld regulates timely patterning transitions during regeneration. A-D)Wg immunostaining in an undamaged disc (A), undamaged disc with Zld inactivated (B), regenerating disc at R48 (C), and regenerating disc at R48 with Zld inactivated (D). E) Zld peak near Wg locus at R48, showcasing wg1_BRV118 region near it. F-K) Bs immunostaining in an undamaged disc (F), undamaged disc with Zld inactivated (G), regenerating disc at R60 (H), regenerating disc at R60 with Zelda inactivated (I), regenerating disc at R72 (J), and regenerating disc at R72 with Zld inactivated (K). Scale Bar: 100µM for all discs.

### Loss of Zld delays the transition to normal patterning in regenerating wing discs

We wondered if Zld’s regulation of targets during late regeneration is an integral aspect of the transition from regeneration back to normal development (Figure 1C-F, R-U). To elucidate the extent to which Zld is important for facilitating timely transitions during late regeneration, we looked at the patterning genes Wg and Bs, which are expressed in unusual patterns during regeneration and return to their original state as the wing disc exits regeneration (Figure 1H-K, W-Z), and can serve as readouts for the re-establishment of many cell fates and patterning elements. Thus, we used Wg and Bs expression to assess the timing of the transitions that occur at the end of regeneration.

To determine the extent to which Zld is important for the late-regeneration transition to the normal Wg expression pattern (Figure 1Y), we examined Wg expression while inactivating Zld. During normal development, inactivation of Zld did not alter Wg expression (Figure 5A-B). During regeneration of control discs at R48, Wg expression was transitioning to its normal expression at the DV boundary and the inner and outer hinge rings in all but 15.3% (n=26) of discs (Figure 5C). However, when we inactivated Zld during regeneration, 48% (n=23) of wing discs retained Wg expression throughout the pouch at R48, appearing similar to R24 wing discs (Figures 1X, 5D). Thus, reducing Zld compromised the timely transition of Wg expression during late regeneration in a significant percentage of regenerating discs.

We wondered if this prolonged ubiquitous expression of Wg in regenerating discs upon Zld inactivation had any impact on regenerative growth. Interestingly, Zld inactivation resulted in a higher percentage of fully regenerated wings (75-100% wing size) (Figure S4A-E), although they had many patterning defects. However, the regenerating wing pouches that lacked Zld function were already larger than controls by R48 (Figure S4F-J), which we speculate could be due to an observed increase in mitoses at R24 but not at R48 (Figure S4K-O). Given that mitoses were not increased at R48, the sustained Wg expression at R48 in discs with inactivated Zld did not appear to enhance proliferation in late regeneration. Interestingly, Zld may reduce proliferation during early wing disc regeneration, and identification of how Zld restricts regenerative growth will be of interest in future studies.

This shift in the timing of the transition back to the normal expression pattern of Wg induced by loss of Zelda may be due to direct regulation of Wg expression by Zld or may be due to an indirect effect on general patterning. Wg expression during disc development and regeneration is regulated by the *wg^1^* and *BRV118* enhancers in the *wg-wnt6* regulatory region (Gracia-Latorre et al., 2022; Harris et al., 2016). Interestingly, during regeneration Zld was bound to a region in the *wg-wnt6* regulatory locus (chr2L:7319693-7321007) that is outside of the *wg^1^*and *BRV118* enhancer region (Figure 5E). Thus, the effect of Zelda on the change in Wg patterning may be direct, but not by modulating the activity at *wg^1^* and *BRV118*, or may be indirect.

We next assessed the transition of Blistered (Bs) expression as regeneration progresses. Importantly, there were no Zld binding sites near *Bs* at R48, so the effects of loss of Zld on Bs expression reflect the overall state of patterning in the regenerating tissue. Bs was barely expressed in the regenerating pouch at R48 (Figure 1J) but returned to its normal expression at R72 (Figure 1K). Hence, we examined the intermediate timepoint of R60. Inactivating Zld in undamaged wing discs had no effect on Bs expression (Figure 5F-G). In control R60 regenerating discs, Bs expression was emerging in the intervein regions, was starting to retreat from the proveins, and was excluded from the D/V boundary in 67% of discs (n=12) (Figure 5H), while the remaining discs had variable Bs expression in the D/V boundary and pro-vein regions. After inactivating Zld, Bs was mis-expressed throughout the regenerating pouch at R60, including in the proveins and the D/V boundary in 73% of discs (n=11) (Figure 5I). While Bs expression had returned to its normal developmental pattern by R72 in 75% of control discs (n=8) (Figure 5J), 57% (n=7) of Zld-inactivated R72 discs had not transitioned correctly, with haphazard Bs expression, especially along the D/V boundary of the wing disc (Figure 5K). Thus, Zld is crucial for the timely transition to normal developmental gene expression patterns during late regeneration, as assessed by Wg and Bs expression.

### Zld is important for stabilizing posterior cell fate

We have previously shown that the posterior cell fate gene *engrailed* (*en*) is susceptible to misregulation by JNK signaling during regeneration, which induces overexpression followed by autoregulated silencing of *en* (Schuster & Smith-Bolton, 2015). To prevent this mis-regulation, the SERTAD ortholog *taranis* (*tara*) and the SWI/SNF BAP complex, defined by the BAP-specific complex member *osa,* act as protective factors to prevent *en* overexpression (Schuster & Smith-Bolton, 2015; Tian & Smith-Bolton, 2021). While *tara* expression is activated in late regeneration, how its expression is activated and how *osa* activity is regulated are not known. Reduction of either *tara* or *osa* during normal development does not cause adult wing defects (Figure S5A,C,E), but their reduction during regeneration leads to adult wings that have a number of anterior features on the posterior half of the wing, namely vein material on the posterior margin, sensory bristles on the posterior margin, an extra anterior cross-vein on the posterior side, anterior compartment shape in the posterior compartment, and distal costa bristles on the posterior side (Figure S5B,D,F) (Schuster & Smith-Bolton, 2015; Tian & Smith-Bolton, 2021). These defects are collectively called posterior-to-anterior (P-to-A) cell fate defects. Interestingly, we noticed for Ac immunostaining that it would often be mis-expressed in the posterior compartment of regenerating discs during Zld inactivation (Figure 4T,U), signifying P-to-A changes are taking place.

Further analysis of adult wings showed loss of Zld did not affect posterior cell fate during normal development (Figure 6A-B). Adult wings after disc regeneration had occasional P-to-A defects, with sporadic anterior sensory bristles and vein material on the posterior margin, as seen previously (Figure 6C) (Schuster & Smith-Bolton, 2015; Tian & Smith-Bolton, 2021). However, adult wings after regeneration with Zld inactivated or knocked down had a much higher frequency of P-to-A defects, including vein material and sensory bristles on the posterior margin, and extra anterior cross veins on the posterior side (Figures 6D, S5H). However, these wings were also often missing large sections of the posterior margin and were missing many or all veins (Figure 4E-F). Thus, we could not quantitate frequency of sensory bristles and vein material on the posterior margin due to the frequent absence of the posterior margin itself, nor could we quantitate frequency of a second anterior crossvein in wings that often lacked all veins.

**Figure 6:**
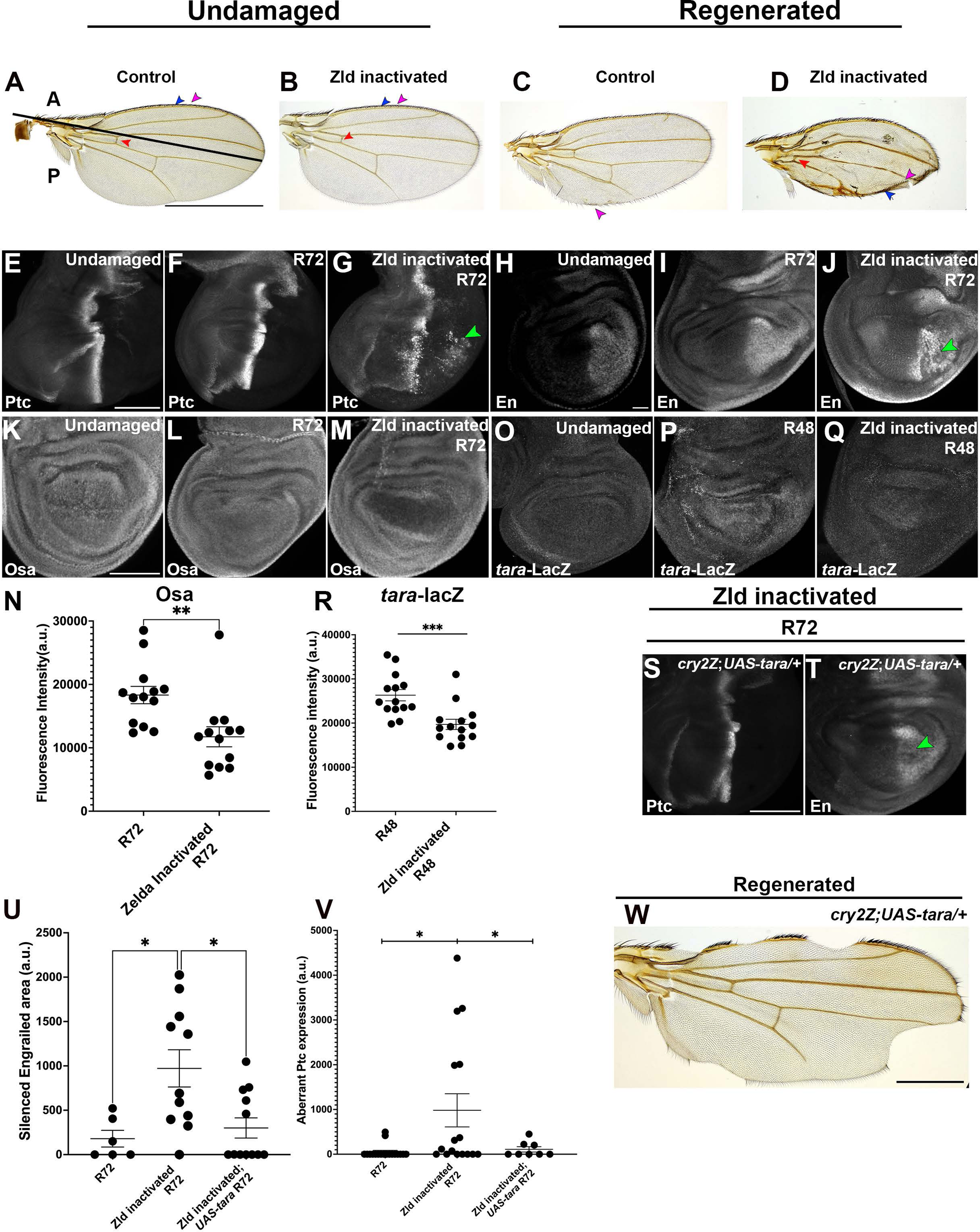
Zld stabilizes posterior cell fate during late regeneration. A-B) Undamaged adult wings, with L1 vein (blue arrowhead), sensory bristles (magenta arrowhead), and anterior crossvein (red arrowhead) marked. Control (A), and with Zld inactivated (B). C-D) Adult wings after disc regeneration. with arrowheads same as in (A), C) control, D) Zld inactivated during regeneration. E-G) Ptc expression in undamaged (E), R72 control (F), and R72 with Zld inactivated (G) wing discs. H-J) En expression in undamaged (H), R72 control (I), and R72 with Zld inactivated (J) wing discs. K-M) Osa expression in an undamaged disc (K), a control R72 disc (L), and an R72 disc that had Zld inactivated (M). N) Quantification of Osa levels in regenerating tissue (n=13 for all comparisons), **p<0.01 Student’s t-test. O-R) *tara-lacZ* enhancer trap expression in an undamaged wing disc (O), an R48 control wing disc (P), and an R48 wing disc with Zld inactivated (Q). R) Quantification of *tara*-lacZ in regenerating tissue (n=14 for all comparisons), ***p<0.001, Student’s t-test. S) Ptc expression in an R72 disc with Tara overexpressed and Zld inactivated. T) En expression in an R72 disc with Tara overexpressed and Zld inactivated. U) Adult wing from regenerated wing disc with Tara overexpressed and Zld inactivated. V) Quantification for En silencing in control R72 discs (n=6), Zld inactivated R72 discs (n=11) and Zld inactivated R7 discs with Tara overexpression (n=11). *p<0.05 One-way ANOVA. X) Quantification of aberrant Ptc expression in control R72 discs (n=18), Zld inactivated R72 discs (n=16) and Zld inactivated R72 discs with Tara overexpression (n=8). *p<0.05 One-way ANOVA. Scale bar for adult wings: 750µM except for Figure 6W (500 µM);. Scale bar for all discs: 100µM.

To confirm that the phenotypes observed in the adult wings were due to P-to-A changes during disc regeneration, we immunostained for the Hedgehog receptor Patched (Ptc). Ptc is expressed in the anterior cells near the anterior-posterior boundary, and during regeneration that expression remained restricted to the anterior side of the anterior-posterior boundary (Figure 6E-F). By contrast, R72 discs that lacked Zld function had Ptc expression in the posterior compartment (Figure 6G), indicating posterior cells that were attaining anterior fate. We also immunostained for the posterior selector transcription factor En, whose misregulation during regeneration is enhanced when levels of the protective factors *tara* or *osa* are reduced (Schuster & Smith-Bolton, 2015; Tian & Smith-Bolton, 2021) (Figure S5I-P). Similarly, when Zld was inactivated in regenerating discs, *en* expression was silenced in portions of the posterior compartment (Figure 6H-J). Thus, Zld plays a role in regulating *en* expression during late regeneration to preserve posterior fate.

We wondered if Zld could help stabilize *en* expression by regulating the expression of *tara* or *osa*. Importantly, our CUT&RUN analysis showed that Zld binds to sites near *en*, *tara* and *osa* in regenerating discs (Figure S5Q,U), raising the possibility of direct and/or indirect regulation of *en* by Zld. Osa protein levels and transcription of *tara* were not altered when Zld was inactivated during normal development (Figure S5 R-T, V-X). In regenerating wing discs, Osa levels were not significantly different between undamaged and control R72 discs (Figure 6K-L).

Interestingly, there was a striking decrease in Osa levels in the regenerating pouch of R72 discs after inactivation of Zld, (Figure 6L-N). In contrast to *osa*, *tara* expression was significantly upregulated in late regenerating discs (Figure 6O-P) (Schuster & Smith-Bolton, 2015). Upon Zld inactivation in R48 discs, *tara* transcription as assessed by an enhancer trap was significantly decreased (Figure 6P-R). Thus, our results suggest Zld can regulate posterior fate in part by controlling expression of the protective factors *osa* and *tara*, to prevent *en* from being silenced during late regeneration.

To confirm these findings, we tried to rescue the P-to-A defects caused by loss of Zld by overexpressing *tara*. Tara overexpression eliminated the ectopic Ptc expression in the posterior of the disc, and largely rescued loss of En in the posterior of the disc, although some discs at R72 still had small areas of En silencing (Figure 6S-V). The resulting adult wings had few P-to-A defects, although additional phenotypes such as loss of margin remained (Figure 6W). Thus, Zld regulates *tara* and *osa*, which prevent loss of posterior fate in late regeneration.

### Zld is important for the structural integrity of the adult wing after disc regeneration

Inactivation of Zld during regeneration led to a high frequency of blisters in the adult wings. There were no blisters after normal development during which Zld was inactivated or knocked down (Figure 7A-B, D). After wing disc regeneration in control animals, about 5% of adult wings had blisters (Figure 7C). Control *pdm2-LexA/+; attP2/+* regenerated discs also regenerated well, with blisters in only 3% of adult wings after disc regeneration (Figure 7D). By contrast, after Zld inactivation during disc regeneration, 23% of the adult wings had blisters (Figure 7E,G-H). RNAi-mediated *zld* knockdown during disc regeneration also caused blisters in 33% of adult wings (Figure 7F,H). Thus, Zld likely controls expression of genes important for the integrity of the adult wing.

**Figure 7:**
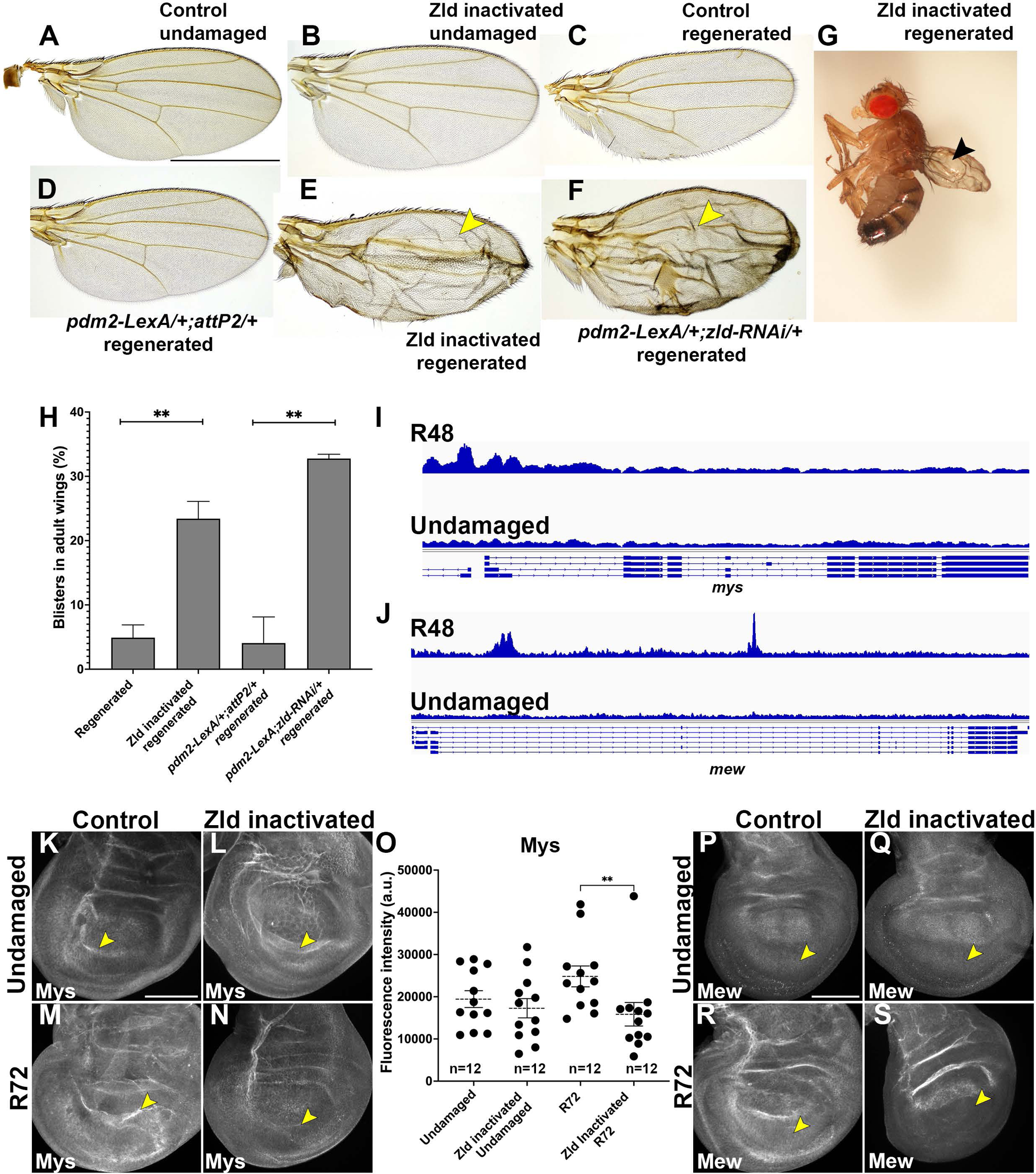
Zld prevents blisters in the adult wing after disc regeneration. A-B) Normal adult wing, control (A) and Zld inactivated (B). C-D) Adult wing after regeneration of a control disc (C) and from *pdm2-LexA/+;attP2/+* control disc (D). E) Adult fly after disc regeneration with Zld inactivated, yellow arrowhead indicates blister. F) Adult wing after disc regeneration with RNAi-mediated *zld* knockdown, yellow arrowhead indicates blister. G) Three-dimensional image of a fly with regenerated adult wing after larval damage with Zld inactivated. Black arrowhead indicates blistered wing. H) Quantification of blisters in adult wings in control (n=207), Zld inactivated (n=190), *pdm2-LexA;attP2/+* RNAi control (n=131) and pdm2-LexA;zld-RNAi/+ (n=91) wings after disc regeneration, **p<0.01, Student’s t-test. I-J) Regions bound by Zld near *mys* (I) and *mew* (J) from CUT&RUN analysis. K-L) Mys expression in undamaged wing discs, with elevated Mys at the D/V boundary (yellow arrowhead). Control (K) and Zld inactivated (L). M-N) Mys expression in R72 discs with yellow arrowhead pointing to the D/V boundary. Control (M), Zld inactivated (N). O) Quantification of Mys expression, **p<0.01, One-way ANOVA. P-Q) Mew expression in wing discs, with yellow arrowhead indicating expression in the ventral half. Undamaged control (P), undamaged with Zld inactivated (Q), R72 control (R), R72 with Zld inactivated (S). Scale bar for adult wings: 750µM. Scale bar for all discs: 100µM.

Integrins play an important role in adhesion of the apposed basal layers of the wing disc when undergoing metamorphosis, and keep the junctions connected as the wing goes through expansion to unfold the adult wing blade after eclosion (Brower et al., 1984; Diaz de la Loza & Thompson, 2017; Fristrom et al., 1993). Interestingly, Zld was bound near the βPS integrin *myospheroid* (*mys*) and αPS1integrin *multiple edamatous wings* (*mew*) at R48.

During normal development, Mys is expressed throughout the wing disc, with slightly elevated levels in the ventral and dorsal regions of the pouch, and higher expression at the D/V boundary (Figure 7K). Inactivation of Zld did not affect Mys expression during normal wing development (Figure 7L). During regeneration, control R72 discs had elevated expression of Mys compared to normal development, especially at the D/V boundary (figure 7M). By contrast, R72 discs in which Zld had been inactivated had reduced expression of Mys with a marked decrease at the DV boundary (Figure 7N-O).

During normal wing development, Mew is expressed in the dorsal and ventral parts of the wing pouch (Figure 7P) and was not affected upon Zld inactivation (Figure 7Q). During regeneration, control R72 discs had high expression of Mew, particularly in the ventral part of the pouch, which was strikingly lost upon Zld inactivation in 68% of R72 discs (n=19) (Figure 7R-S). As expression of integrins in both dorsal and ventral halves of the disc is crucial for maintaining junctions and cell adhesion while the disc undergoes apposition and everts, loss of Mys at the D/V boundary and loss of Mew in the ventral half of the disc likely contribute to the observed blisters.

## Discussion

Here we have shown that the pioneer transcription factor Zld is crucial for cell fate, patterning, and integrin gene expression as the regenerating wing disc transitions to normal development during late regeneration. Regeneration has long been thought to deploy developmental mechanisms to repair and regrow the tissue (Iismaa et al., 2018; N. Suzuki & Ochi, 2020), a notion that has been challenged recently. For example, amniote limb regeneration studies have shown that *fgf10*, a gene important for normal limb development, is not important for limb regeneration (M. Suzuki et al., 2024). Conversely, *tara* and *brat* are required for cell fate preservation in regenerating *Drosophila* wing imaginal discs but not in normally developing discs (Abidi et al., 2023; Schuster & Smith-Bolton, 2015). Moreover, there is recent evidence of regeneration-specific factors that initiate developmental processes in tissues undergoing regeneration after the initial damage response. For instance, during Xenopus limb regeneration, transcription factors Hox12/Hoxc13 promote expression of pivotal developmental genes, while being dispensable for typical limb development (Kawasumi-Kita et al., 2024). In *Drosophila* wing discs, Zld appears to have an analogous role, as it is critical for re-instating the developmental program during late regeneration but is not required for normal development. Intriguingly, one report shows overexpressing Zld during normal development results in blisters in the adult wing (Staudt et al., 2006), suggesting Zld may be capable of regulating integrins when overexpressed. However, in the case of regeneration, it is loss of Zld that causes reduction in integrin expression in the disc and blisters in the adult wing, not overexpression.

While Zld is an established pioneering transcription factor in the *Drosophila* embryo through its recognition of CAGGTA sites (Harrison et al., 2011), Zld occupancy of these sites in the neuroblast (Larson et al., 2021) is not as enriched, nor are they occupied by Zld in the regenerating wing disc (Figure 3C). Furthermore, there is little overlap between Zld binding sites in the neuroblast and the regenerating wing disc, nor are there common potential cofactors based on motif predictions, except for Trl/CLAMP. Thus, it is unclear whether Zld acts as a pioneer factor during disc regeneration or what directs tissue-specific binding of Zld. We did find enrichment for binding motifs for transcription factors such as Fkh and Rib in our CUT&RUN data. These or other transcription factors could help direct Zelda binding during late regeneration.

In addition to regulating target genes directly, we established that Zld can influence cell fate through Notch signaling, as we confirmed that the Notch ligand Dl is a target of Zld (Figure S3E,F-H, Figure 4N). Interestingly, we also found Zld bound near additional Notch signaling pathway components and regulators, including *E(spl)Mβ*, *serrate*, *notch,* and *mastermind*. Thus, Zld could be regulating Notch signaling in general, exacerbating some of the Zld phenotypes related to *ct* and *ac* as they are downstream of Notch as well as direct targets of Zld.

A recent report demonstrated that Zld is important for appendage regeneration in the American cockroach (Zhang et al., 2024), establishing Zld as a regeneration factor in a different insect and context. In another invertebrate, the acoel worm, the *early growth factor* (*egr*) was shown to act as a pioneer factor to regulate early wound genes (Gehrke et al., 2019), reinforcing the importance of pioneer factors during regeneration. While a Zld homolog is not found in vertebrates, vertebrates do have pioneer factors such as Sox2, Klf4, and Oct-4, which may be required for cellular reprogramming during regeneration. Indeed, Sox2 and Klf4 are required for reprogramming retinal cells during retinal ganglion cell regeneration (Rocha-Martins et al., 2019). Thus, understanding the functions of transcription factors like Zld will provide insights into how pioneer factors may play distinctive roles in initiating the transition from a regenerative tissue back to normal development.

## STAR METHODS

### KEY RESOURCES TABLE

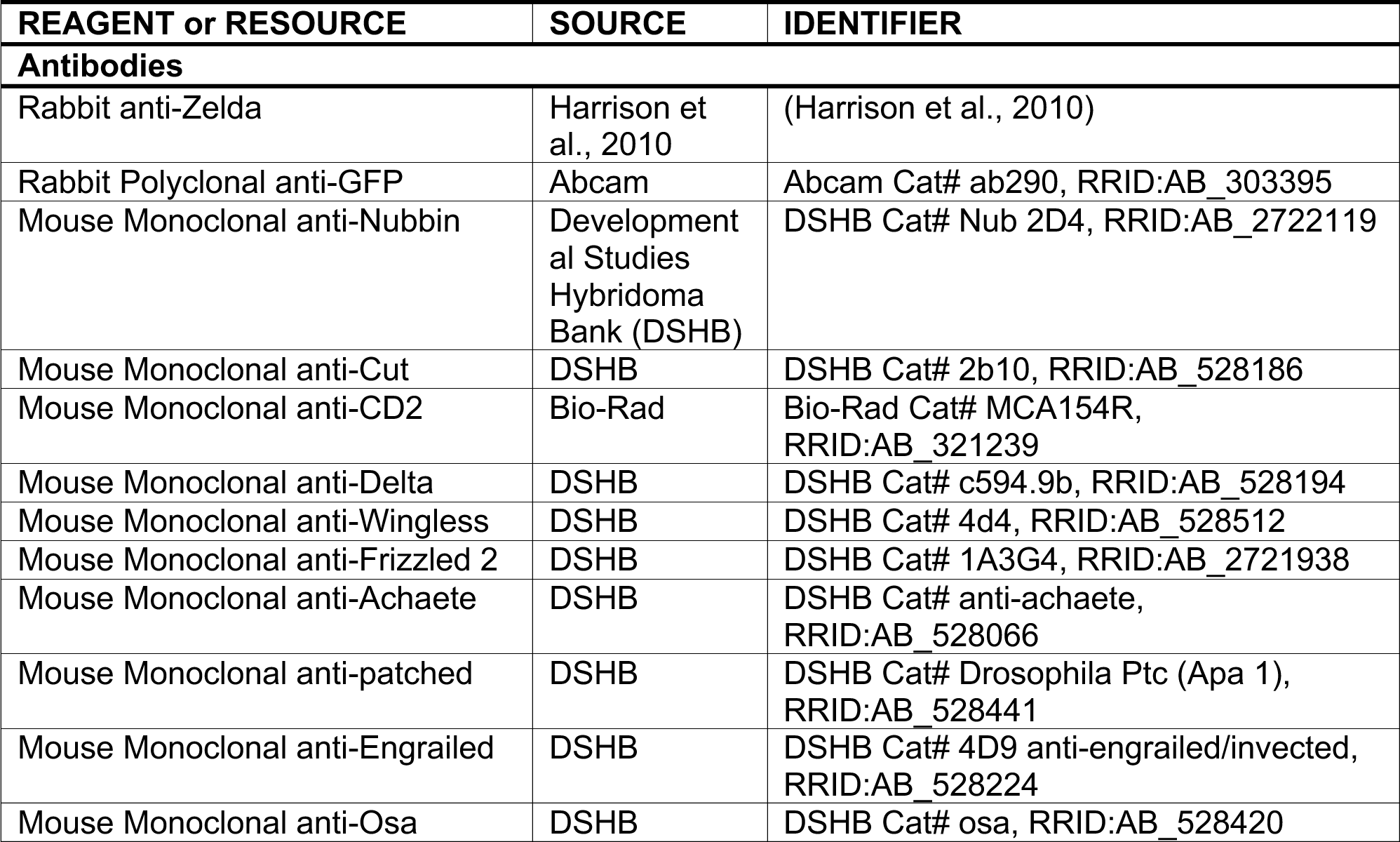
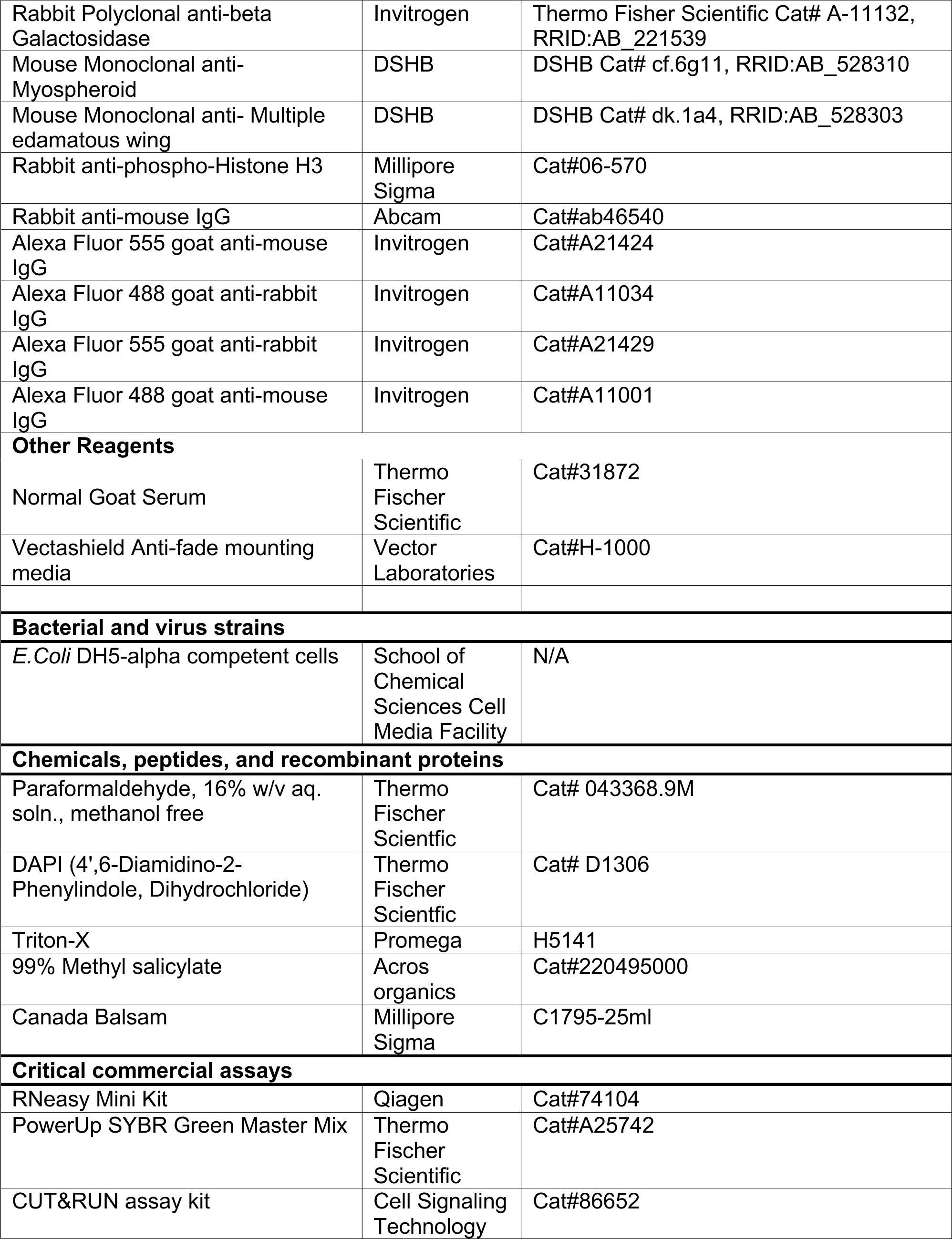
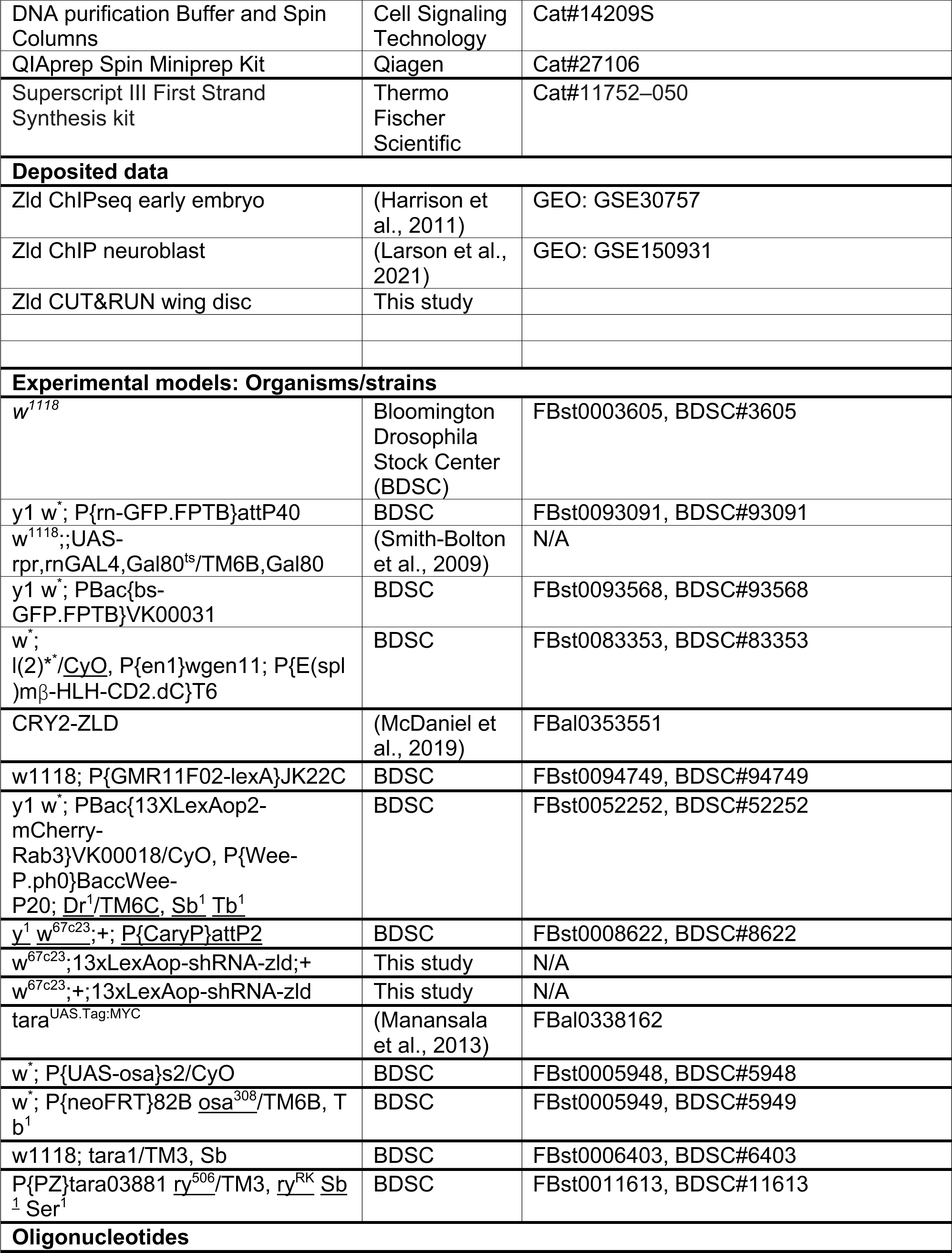
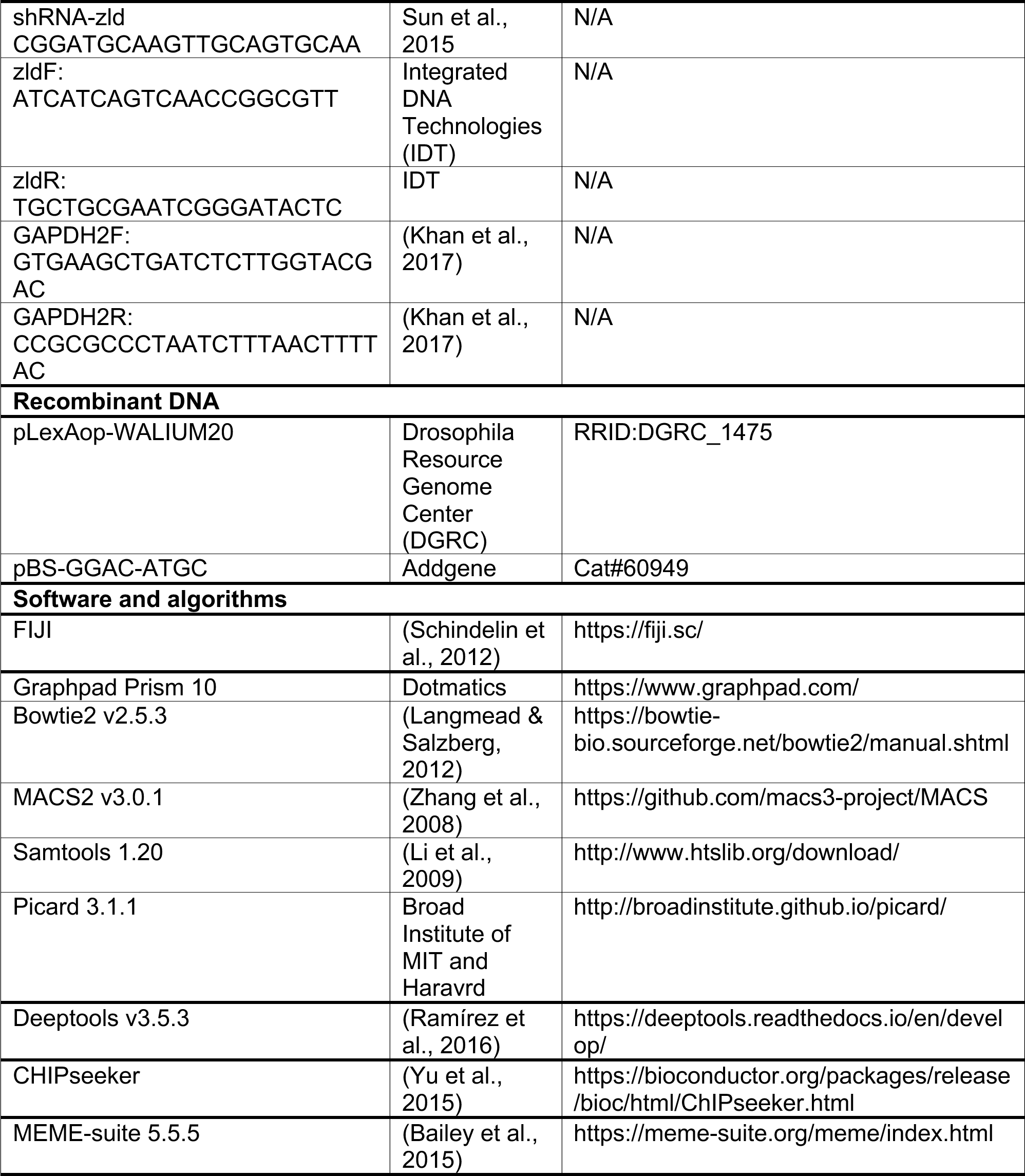
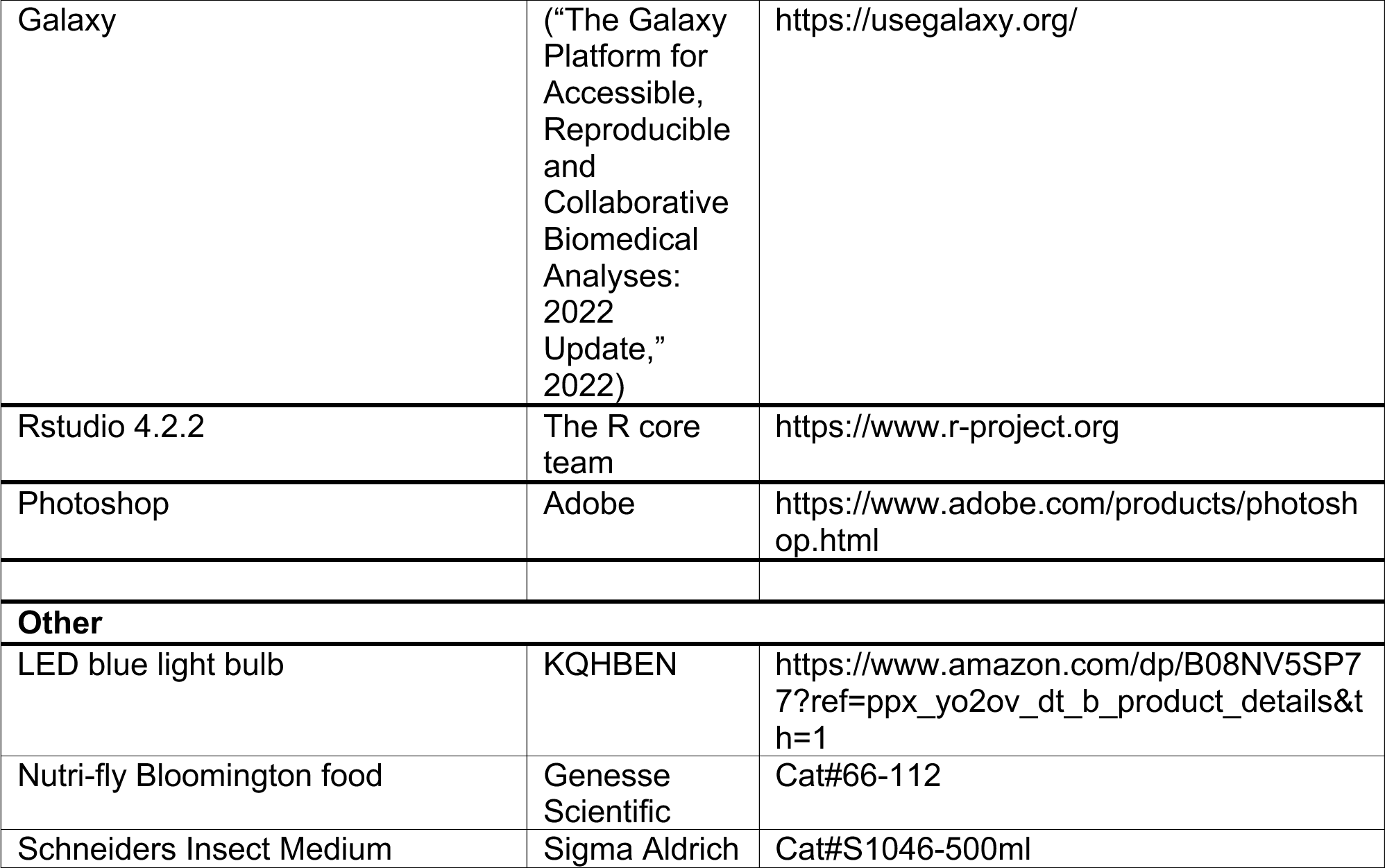

#### Methods

##### Tissue ablation

Ablation and regeneration studies were carried out as previously described (Smith-Bolton et al., 2009). Briefly, egg lays were carried out at room temperature in the dark for 4 hours on grape juice plates with yeast paste, after which they were kept at 18°C. Approximately 50 hatched larvae were collected in separate Nutri-fly Bloomington food vials with yeast paste added and food churned. At early day 7, when larvae are at early third instar stage as determined by mouth-hooks, damage was induced by *reaper* by placing the vials in a circulating water bath at 30°C for 24 hours. The vials were then cooled in an ice-water bath for 60 seconds and returned to an 18°C incubator. The regenerating animals were either dissected according to the required time point or were allowed to develop to adulthood to assess adult wing size and patterning. Undamaged animals were kept consistently at 18°C and were dissected during their third instar crawling stage.

##### Optogenetic inactivation of Zelda

Optogenetic inactivation experiments were carried out by placing food vials containing undamaged or damaged larvae about 3 inches away from blue LED lights inside an 18°C incubator, thus ensuring temperature was controlled. Blue light exposure occurred between the 8th and 11th days after egg lay for both damaged and undamaged animals homozygous for *CRY2::zld* that had spent 24 hours in the 30°C water bath. For experiments in which Zld was inactivated in flies containing Bs-GFP, UAS-tara and UAS-osa, only males hemizygous for *CRY2::zld* were dissected and/or analyzed. Undamaged animals for these experiments were exposed to equal amount of blue light but kept at 18°C throughout. For control experiments, *CRY2::zld* flies were continuously kept in the dark to ensure no Zld inactivation caused by normal bright light.

##### Immunohistochemistry

Immunostaining was carried out as previously described (Smith-Bolton et al., 2009). Primary antibodies were used at the following concentrations: anti-Nubbin (1:100), anti-Cut (1:150), anti-CD2 (1:500), anti-Delta (1:400), anti-Wingless (1:100), anti-Frizzled2 (1:50), anti-Achaete (1:10), anti-Patched (1:50), anti-Engrailed (1:3), anti-Osa (1:1), anti-βgal (1:500), anti-Myospheroid (1:50), anti-Mew (1:50) and anti-phospho-Histone-H3 (1:500). Secondary antibodies with Alexa fluorophores and DAPI were used at a 1:1000 dilution.

All immunofluorescence images were captured using a Zeiss LSM700 or LSM900 confocal microscope. Maximum intensity projections were used for quantifying protein expression pattern or intensity in the regenerating pouch, which was demarcated by Nubbin expression or by generating a 150×150 pixel area where the presumptive pouch would be. Images were processed and quantified using FIJI, and parameters and settings for quantified images were kept the same for all images in a given experiment. Percent of DV boundary expressing Cut was calculated using the freehand line tool in FIJI to draw a line across Cut expression along the D/V boundary and measuring length of expression divided by full D/V boundary length. Area of En silencing in the posterior compartment was calculated by outlining the area where En was repressed using the freehand selection tool in FIJI. The same method was used for calculating the ectopic Ptc expression in the posterior compartment.

##### Adult wing quantification

Adult wings were mounted in Gary’s Magic mount media (Canada Balsam dissolved in Methyl salicylate). Images were taken on an Echo Revolve R4 microscope and were processed using FIJI. Percent of veins present was calculated using the freehand line tool in FIJI to measure actual vein length divided by presumptive full vein length.

##### qPCR experiment

qPCR experiments were carried out as described previously (Abidi et al., 2023). Briefly, 20-30 discs were collected in Schneiders Insect Medium, and the total RNA was extracted using the RNaeasy mini kit. Approximately 100ng of RNA was used for each sample to make cDNA using the Superscript III First Strand Synthesis kit. The qPCR reaction was made with addition of SYBR green mastermix and then run on an ABI Step One Plus Real Time PCR System. Each experiment included three biological replicates, with three technical replicates each. Fold changes were then calculated according to the ΔΔC_t_ method, using GAPDH2 as a comparative housekeeping gene.

##### CUT&RUN experiment

For the CUT&RUN experiments, we used *zldGFP;+;UAS-rpr,rnGAL4,Gal80^ts^ /+* flies for regenerating discs and *zldGFP;+;+* for undamaged flies. Control experiments also used *w^1118^;+; UAS-rpr,rnGAL4,Gal80^ts^/+* discs, that lacked GFP-tagged Zelda. Two replicates were performed for each genotype and condition. All reagents used were from the CUT&RUN assay kit. For each replicate, 100 wing discs were dissected from R48 or undamaged larvae into 1x wash buffer (350*μL* 10X Wash buffer, 35*μL* 100X Spermidine, 17.5*μL* 200X Protease inhibitor cocktail and 3097.5*μL* Nuclease free water). Discs were then suspended in activated Concanavalin A beads, after which Antibody binding buffer (1*μL* 100X Spermidine, 0.5*μL* 200X Protease inhibitor cocktail, 2.5*μL* Digitonin solution, 0.2*μL* GFP antibody and 95.8*μL* Antibody buffer) was added to the sample for primary antibody binding and kept on a nutator overnight in 4°C. Next, the samples were treated with 1.5*μL* pAG-MNase enzyme in 50*μL* digitonin buffer (4.56*μL* 10X wash buffer, 0.46*μL* 100X spermidine, 0.23*μL* 200X protease inhibitor cocktail, 1.14*μL* digitonin solution and 40.6*μL* nuclease free water), followed by incubation at 4°C for one hour. DNA digestion was performed by activating the pAG-MNase enzyme with 3*μL* CaCl_2_ for 30 minutes at 4°C. The reaction was stopped and chromatin was released by adding the sample with 150*μL* of 1X stop buffer, which included the spike-in yeast DNA (37.5*μL* 4X stop buffer, 3.75*μL* Digitonin solution, 0.75*μL* RNase A, 5*μL* of yeast spike-in and 103*μL* nuclease free water), incubating at 37°C for 15 minutes. The released chromatin was then purified using the DNA purification Buffer and Spin Columns kit.

The libraries were prepared by the Roy J. Carver Biotechnology Center, DNA services facility using the UltraLow Input Library construction kit from Tecan. Samples were sequenced in the DNA Services facility on a NovaSeq 6000 (101 cycles) from both ends using V1.5 sequencing kits. Fastq files were generated and demultiplexed with the bcl2fastq v2.20 Conversion Software (Illumina).

##### CUT&RUN data processing

The CUT&RUN paired-end reads were aligned to the *Drosophila melanogaster* dm6 genome (Release 6 plus ISO1 MT, version 2014) and *Saccharomyces cerevisiae* genome version R64 using bowtie2. Alignment parameters included --very-sensitive-local --no unal –no-mixed --no-discordant --phred33 -I 10 -X 700 –local--no-overlap --no dovetail. Reads were then sorted according to coordinates using Samtools. The resultant files were processed with MarkDuplicates for identifying and removing PCR duplicates with high stringency. The coverage file was normalized and scaled in accordance to reads aligned to the spike-in (*S. cerevsiae*) using Genomcov, part of Bedtools. The continuous coverage file was obtained using Bamcoverage, part of Deeptools, which was normalized either to spike-in DNA or to 1x depth (reads per genome coverage). Data range for coverage files in each figure was set between 0-10 except for *achaete* (0-4.5), *tara* (0-15), and *osa* (0-15). Peak calling was done using MACS2 with default parameters. Peak files were annotated using ChIPseeker, and a heatmap was made using plotHeatmap, part of Deeptools. Peak-called files were converted to FASTA sequence using Extract Genomic DNA tool, part of Galaxy tools. Novel motif discovery and analysis was carried out using peak-called FASTA files in MEME-ChIP in discriminative mode, part of MEME-Suite.

Generated CUT&RUN data was compared to ChIP-seq data for Zelda in stage 14 embryo (GSE 30757 and Type II neuroblast (GSE 150931).

##### LexAop-shRNA-*zld* generation

For making a stable *Drosophila* line with RNAi-mediated knockdown of *zld*, the passenger strand previously described (Sun et al., 2015) specific to the *zld* transcript was cloned into a LexAop-pWALIUM20 vector using services from GenScript. The hairpin sequence used was TAGTTATATTCAAGCATA. The plasmid was then injected into embryos and integrated into the *Drosophila* genome at the attP40 and attP2 recombination sites using services from BestGene.

##### Statistical analysis

For measuring relevant statistical differences of means between two groups, the students t-test method was applied. For comparing multiple groups, the One-way ANOVA statistical test was applied. For comparing wing sizes between different groups, a Chi-squared test was performed. Analysis and graphs were made using Graphpad Prism 10.

## Data availability

Sequencing data for the CUT&RUN experiments are available from the GEO database, accession number GSE268674. All other relevant data are available at the Illinois Databank (insert URL here prior to publication) and upon request.

## Author contributions

A.B. designed, interpreted, and carried out the experiments in this study. K.S. designed and carried out preliminary studies. C.K. carried out and analysed data from experiments related to adut wing defects and *engrailed* expression. S.S. optimized the CUT&RUN protocol. R.K.S.-B. designed and interpreted experiments. The manuscript was written by A.B. and R.K.S.-B, with suggested edits by K.S. and S.S.

## Declaration of interests

The authors declare no competing interests. K.S.’s current institutional affiliation is the Center for Developmental Genetics, New York University. S.S.’s current institutional affiliation is the Roy J. Carver Biotechnology Center, University of Illinois at Urbana-Champaign.

## Acknowledgements

We would like to thank Snigdha Mathure and Felicity Hsu for critical reading of the manuscript. We would like to thank Dr. Melissa Harrison for providing many of the reagents used in this study. We would like to give a special thanks to BDSC, DGRC and DSHB for reagents. This work was supported by NIH grants R01GM107140 and R35GM14174, and an Arnold O. Beckman Award from the University of Illinois (RB20123).

---

**Figure S1:**
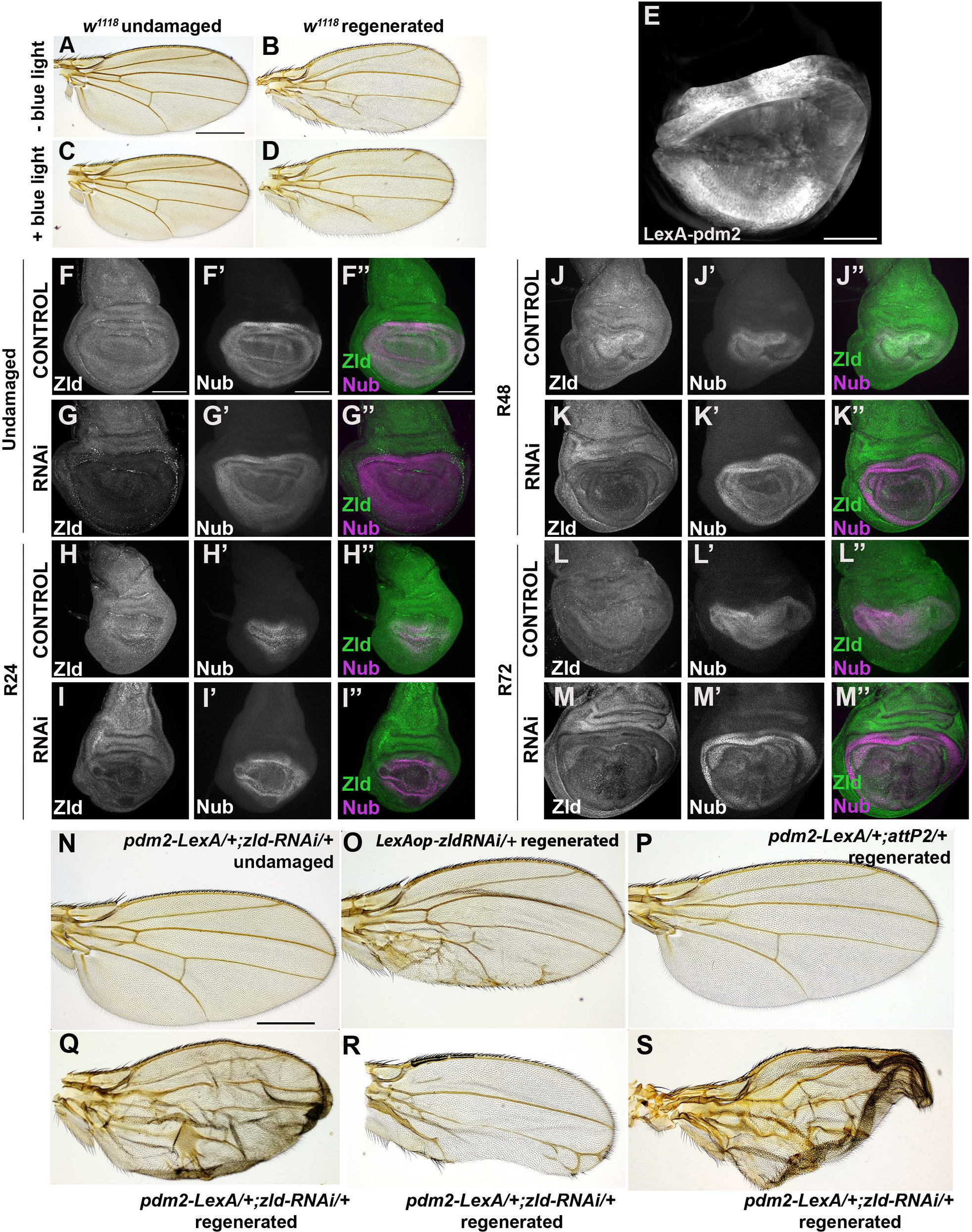
RNAi-mediated Zld knockdown in regenerating wing discs causes adult wing defects. A-B) Adult wings from animal raised without blue light. A) Undamaged adult wing. B) Adult wing from regenerated wing disc. C-D) Adult wings from animal raised with blue light. C) Undamaged adult wing. D) Adult wing from regenerated wing disc. E) *LexAop* transgene expressing 6x-mcherry driven by *pdm2-lexA*. F-M) Zld and Nub co-immunostaining in control and *zldRNAi* discs when undamaged (F-G”) and at regeneration time points R24 (H-I”), R48 (J-K”) and R72 (L-M”). N) Undamaged adult wing after *zld* knockdown. O) Adult wing from *LexAop-zldRNAi* regenerated disc. P) Adult wing from *pdm2-LexA/+;attP2/+* regenerated disc. Q-S) Adult wings after *zld* knockdown with *pdm2-LexA/+; LexAop-zldRNAi /+* during regeneration. Scale bar for adult wings: 500µM. Scale bar for all discs: 100µM.

**Figure S2:**
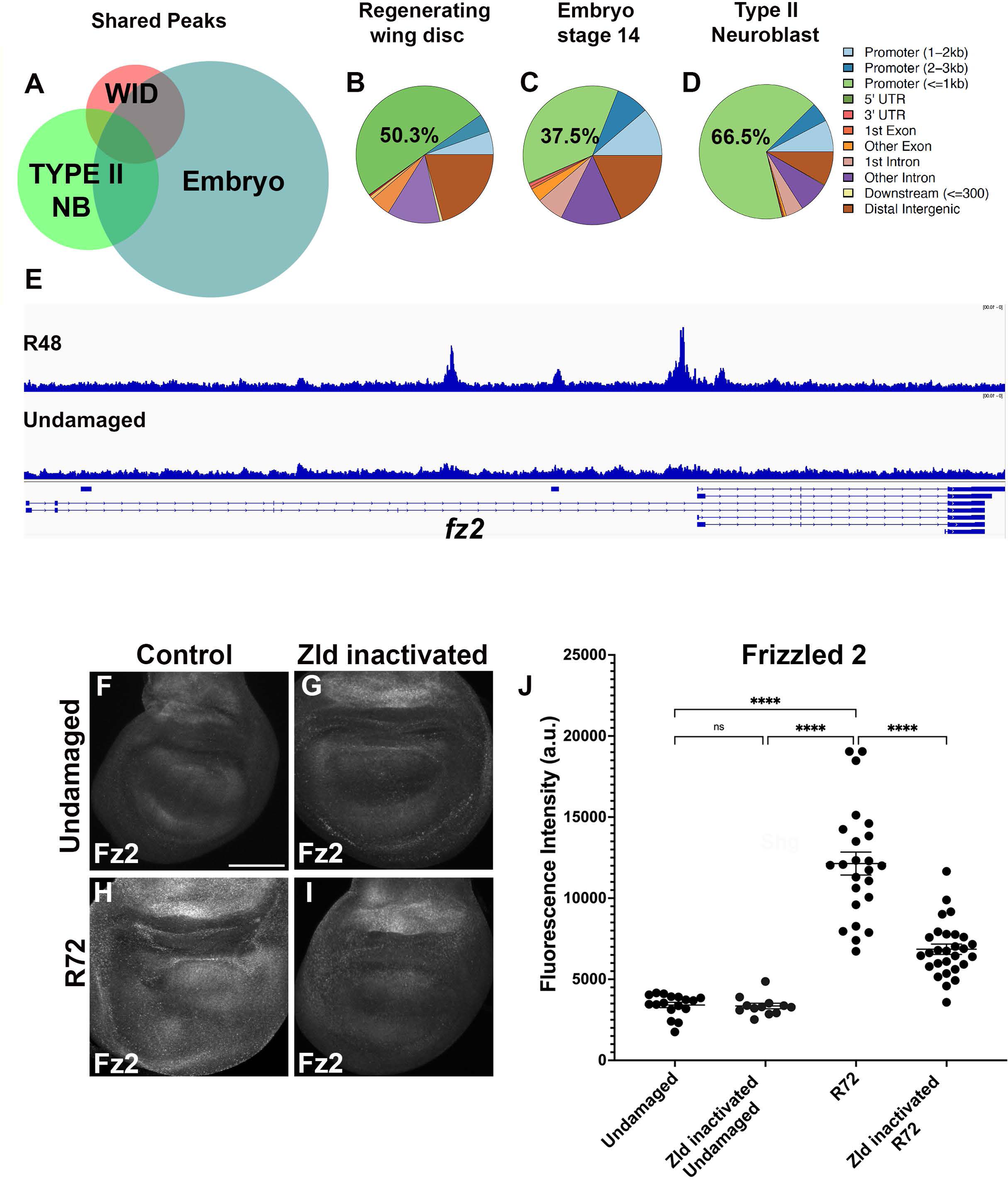
Comparison of Zld binding sites among tissues, and validation of a putative target. A) Shared Zld binding sites among wing imaginal disc late regeneration (WID)(3541 peaks), Stage 14 embryo (20,119 peaks) and Type II neuroblasts (NB) (7215 peaks). Overlap between the late regenerating wing disc and embryo is 2449 peaks, between the late regenerating wing disc and neuroblast is 1105 peaks, and between the neuroblast and embryo is 2865 peaks. B-D) Distribution of genomic location of annotated peaks using ChIPseeker in late wing disc regeneration (B), stage 14 embryos (C), and type II neuroblasts (D). E) Zld binding in R48 regenerating discs near the gene *fz2*. F-I) Fz2 expression in an undamaged wing disc (F), undamaged wing disc with Zld inactivated (G), R72 wing disc (H), and R72 wing disc with Zld inactivated (I). J) Quantification of Fz2 expression in undamaged control (n=17), undamaged Zld inactivated (n=12), R72 control (n=24), and R72 Zld inactivated (n=28) discs. Scale bar: 100µM for all discs.

**Figure S3:**
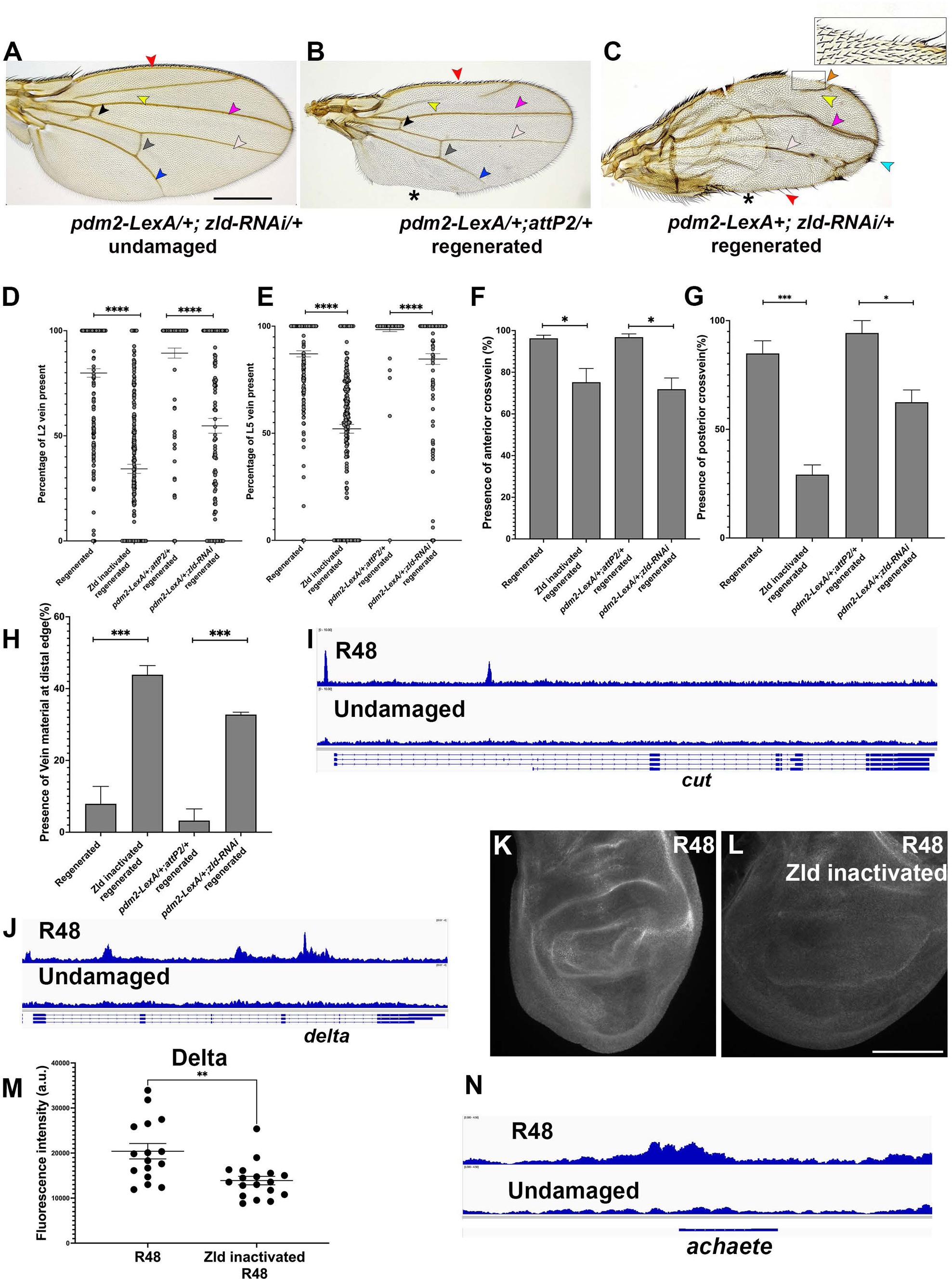
Zld is important for sensory bristle and pro-vein identity in regenerating wing discs. A) Undamaged adult wing from animal with Zld knocked down during development showing L1 (red arrowhead), L2 (yellow arrowhead), L3 (magenta arrowhead), L4 (white arrowhead) and L5 (blue arrowhead) veins. B) Adult wing after disc regeneration, with same arrowheads as in (A). * marks missing posterior margin. C) Adult wing after disc regeneration with Zld knocked down by RNAi, arrowheads same as in B, along with missing sensory bristles (orange arrowhead) and extra vein material on distal edge (teal arrowhead). D-H) Quantification of percentage of L2 vein present (D), percentage of L5 vein present (E), presence of ACV (F), presence of PCV (G), presence of excess distal vein material (H), in control (n=207), Zld inactivated (n=190), *pdm2-LexA;attP2/+* (n=131) and pdm2-LexA; zld-RNAi/+ (n=91) wings after disc regeneration. *p<0.05, ***p<0.001, ****p<0.0001, Student’s t-test. I) Zld-occupied sites at the promoter region of *ct* at R48. J) Zld-occupied sites at the at the *Dl* locus at R48. K-L) Dl expression in regenerating wing disc at R48 (K), and regenerating wing disc at R48 with Zld inactivated (L). (M) Quantification of Dl expression, **p<0.01 (Students t-test). N) Zld-occupied site at the *achaete* locus at R48. Scale bar for adult wings: 500µM. Scale bar for all discs: 100µM.

**Figure S4:**
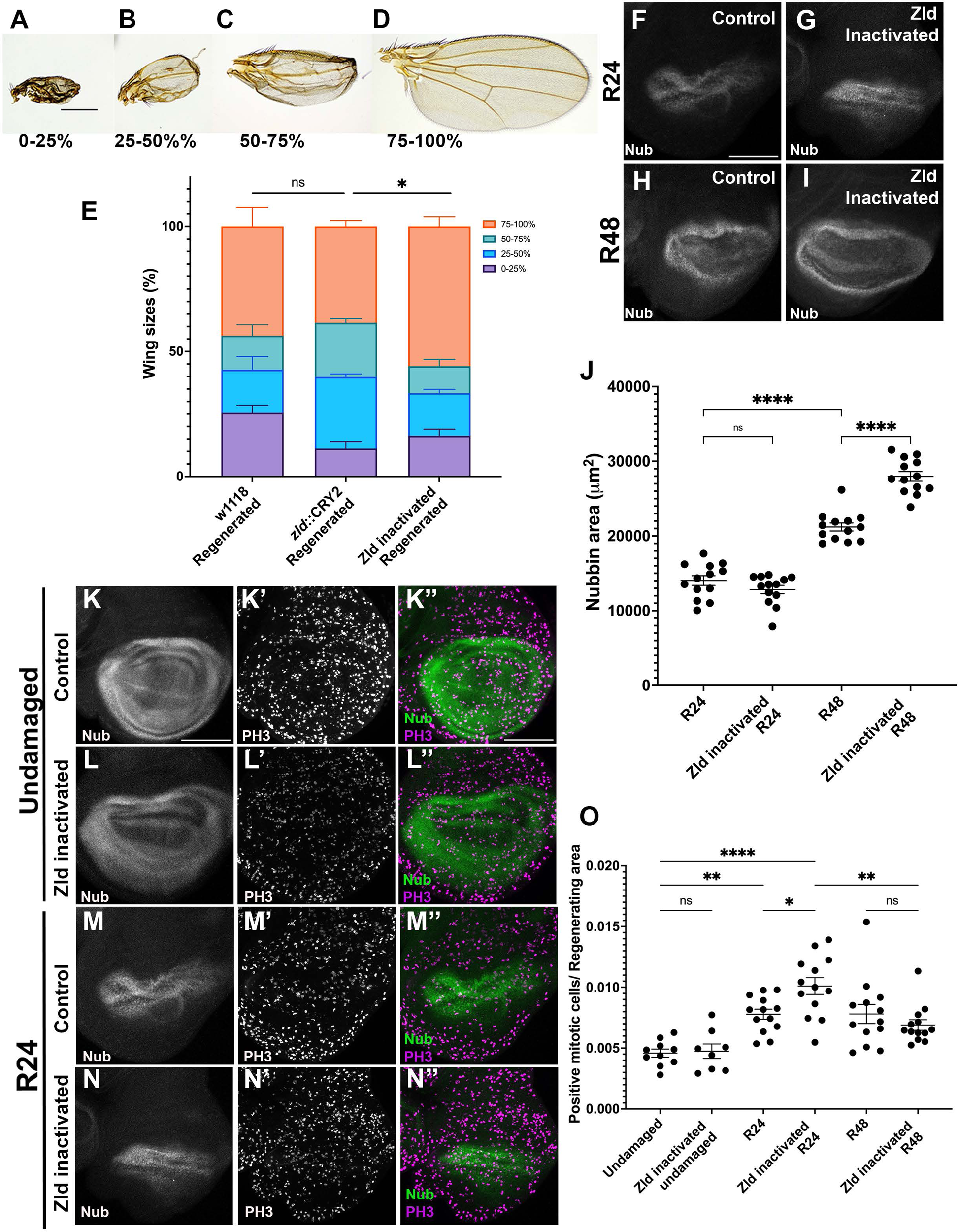
Zld restricts proliferation in wing disc regeneration. A-D) Adult wings categorized according to size compared to undamaged adult wing. 0-25% sized wing (A), 25-50% sized wing (B), 50-75% sized wing (C), 75-100% sized wing (D). E) Adult wing sizes for *w^1118^;+;UAS-rpr,rnGAL4,Gal80^ts^/+* regenerated (n=258), *zld::cry2;+;UAS-rpr,rnGAL4,Gal80^ts^/+* regenerated without blue light (n=387), and *zld::cry2;+;UAS-rpr,rnGAL4,Gal80^ts^/+* regenerated with blue light (n= 446). 0-25% (purple), 25-50% (blue), 50-75% (teal), 75-100% (orange). **p<0.01, Chi-square test. F-I) Nub immunostaining in R24 and R48 discs for *zld::cry2;+;UAS-rpr,rnGAL4,Gal80^ts^/+* regenerating flies. (F) Control R24 disc (no blue light), (G) R24 disc with Zld inactivated, (H) Control R48 regenerating disc (no blue light) and I) R48 wing disc with Zld inactivated. J) Quantification of regenerating pouch area demarcated by Nubbin in R24 and R48 regenerating discs (n=13 for all comparisons), ns p>0.05, *p<0.05, ****P<0.0001, Student’s t-test. K-L”) Nub (K,L), PH3 (K’,L’) staining and composite image (K’’,L’’) in undamaged control discs without blue light and undamaged discs with Zld inactivated (L-L’’’). M-N”) Nub (M,N) and PH3 (M’,N’) staining and composite image (M’’,N’’) in R24 regenerating control discs without blue light (M-M’’) and discs with Zld inactivated (N’N’’). O) Quantification of number of mitotic cells per Nub-expressing area at R24 and R48 (n=13 for all comparisons), ns P> 0.5, *P<0.05, **P<0.01, ***P<0.001, ****P<0.0001, One-way ANOVA test. Scale bar: 500 µM for all adult wings. Scale bar: 100µM for all discs.

**Fig S5:**
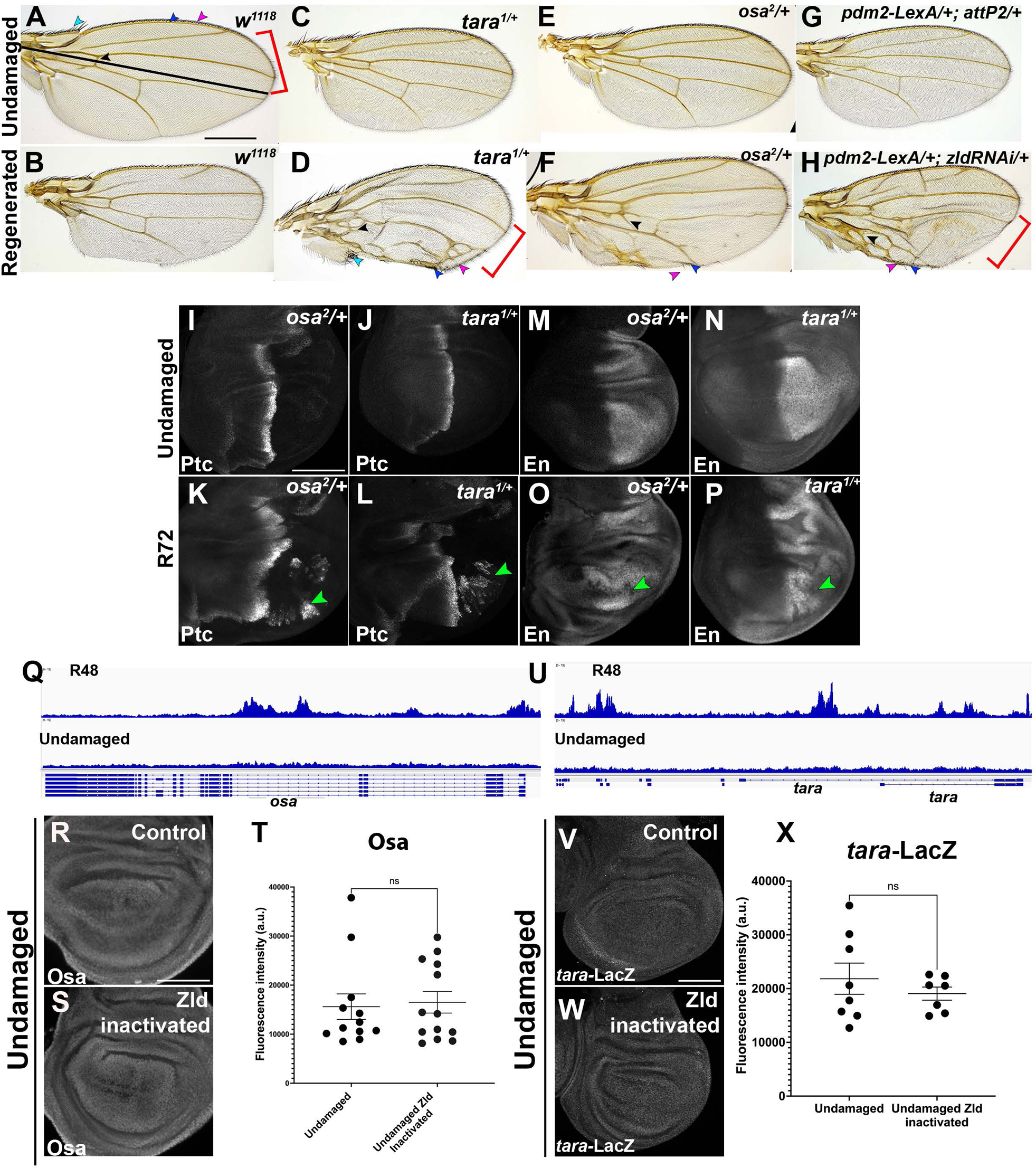
Zld is important for posterior cell fate during regeneration. A) *w^1118^* adult wing showing anterior L1 vein (blue arrowhead), anterior sensory bristles (magenta arrowhead), anterior compartment shape (red bracket) and anterior crossvein (black arrowhead). B) Adult wing from *w^1118^* regenerated disc. C) Undamaged *tara^1^/+* adult wing. D) Adult wing from *tara^1^/+* regenerated disc showing anterior features as in (A) in posterior of wing. E) Undamaged *osa^2^/+* adult wing. F) Adult wing from *osa^2^/+* regenerated disc showing anterior features as in (A) in the posterior of the wing. G) Adult wing from *pdm2-LexA /+;attP2/+* regenerated disc. H) Adult wing from *pdm2LexA; zldRNAi/+* regenerated disc showing anterior features as in (A) in the posterior of the wing. I-L) Ptc expression in *osa^2^/+* undamaged wing disc (I), *tara^1^/+* undamaged wing disc (J), *osa^2^/+* R72 wing disc (K), and *tara^1^/+* R72 wing disc (L). Green arrowheads show aberrant Ptc expression. M-P) En expression *osa^2^/+* undamaged wing disc (M), *tara^1^/+* undamaged wing disc (N), *osa^2^/+* R72 wing disc (O), and *tara^1^/+* R72 wing disc (P). Green arrowheads show En silencing. Q) Zld bound near the *osa* locus in R48 regenerating discs and in undamaged discs R) Osa expression in undamaged wing disc. S) Osa expression in undamaged disc with Zld inactivated. T) Quantification of Osa immunostaining in control undamaged (n=12) and Zld inactivated undamaged (n=13) discs, ns p>0.5, Student’s t-test. U) Zld bound near the *tara* locus in R48 regenerating discs and in undamaged discs. V) *tara-lacZ* expression (beta-galactosidase immunostaining) in undamaged wing disc. W) *tara-lacZ* expression in undamaged disc with Zld inactivated. X) Quantification of β-galactosidase in control undamaged (n=8) and Zld inactivated undamaged (n=7) discs, ns p>0.5, Student’s t-test. Scale bar for adult wings: 500µM. Scale bar for all discs: 100µM.

## Notes

### Competing Interest Statement

The authors have declared no competing interest.

